# Deconvolution of Transcriptional Networks Identifies TCF4 as a Master Regulator in Schizophrenia

**DOI:** 10.1101/133363

**Authors:** Abolfazl Doostparast Torshizi, Chris Armoskus, Hanwen Zhang, Marc P. Forrest, Siwei Zhang, Tade Souaiaia, Oleg V. Evgrafov, James A. Knowles, Jubao Duan, Kai Wang

**Affiliations:** Raymond G. Perelman Center for Cellular and Molecular Therapeutics, Children’s Hospital of Philadelphia, Philadelphia, PA 19104, USA; Department of Pathology and Laboratory Medicine, Perelman School of Medicine, University of Pennsylvania, Philadelphia, PA 19104, USA; College of Medicine, SUNY Downstate Medical Center, Brooklyn, NY 11203, USA; Zilkhe Neurogenetic Institute, University of Southern California, Los Angeles, CA 90089, USA; Center for Psychiatric Genetics, North Shore University Health System, Evanston, IL 60201, USA; Department of Physiology, Feinberg School of Medicine, Northwestern University, Chicago, IL 60611, USA; Center for Autism and Neurodevelopment, Feinberg School of Medicine, Northwestern University, Chicago, IL 60611, USA; Department of Psychiatry and Behavioral Neurosciences, University of Chicago, Chicago, IL 60015, USA

## Abstract

Tissue-specific reverse engineering of transcriptional networks has uncovered master regulators (MRs) of cellular networks in various cancers, yet the application of this method to neuropsychiatric disorders is largely unexplored. Here, using RNA-Seq data on postmortem dorsolateral prefrontal cortex (DLPFC) from schizophrenia (SCZ) patients and control subjects, we deconvolved the transcriptional network to identify MRs that mediate expression of a large body of target genes. Together with an independent RNA-Seq data on cultured cells derived from olfactory neuroepithelium, we identified *TCF4*, a leading SCZ risk locus implicated by genome-wide association studies, as one of the top candidate MRs that may be potentially dysregulated in SCZ. We validated the dysregulated TCF4-related transcriptional network through examining the transcription factor binding footprints inferred from human induced pluripotent stem cell (hiPSC)-derived neuronal ATAC-Seq data, as well as direct binding sites obtained from ChIP-seq data in SH-SY5Y cells. The predicted *TCF4* transcriptional targets were enriched for genes showing transcriptomic changes upon knockdown of *TCF4* in hiPSC-derived neural progenitor cells (NPC) and glutamatergic neurons (Glut_N), based on observations from three separate cell lines. The altered *TCF4* gene network perturbations in NPC, as compared to that in Glut_N, was more similar to the expression differences in the *TCF4* gene network observed in the DLPFC of individuals with SCZ. Moreover, *TCF4*-associated gene expression changes in NPC were more enriched than Glut_N for pathways involved in neuronal activity, genome-wide significant SCZ risk genes, and SCZ-associated *de novo* mutations. Our results suggest that *TCF4* may potentially serve as a MR of a gene network that confers susceptibility to SCZ at early stage of neurodevelopment, highlighting the importance of network dysregulation involving core genes and many hundreds of peripheral genes in conferring susceptibility to neuropsychiatric diseases.

## Introduction

Schizophrenia (SCZ) is a debilitating neuropsychiatric disease, which affects about 1% of adults (*1, 2*). SCZ is a highly heritable disease, with heritability estimated to be up to 73%-83% in twin studies (*3*); however, its underlying genetic architecture remains incompletely understood (*4*). Recent genome wide approaches have identified a plethora of common disease risk variants (*5*), rare copy-number variants (CNVs) of high penetrance (*6, 7*) and rare protein-altering mutations (*8*) that confer susceptibility to SCZ. Although exciting, the biology underlying these genetic findings remains poorly understood, prohibiting the development of targeted therapeutic strategies. A major challenge is the polygenic nature of SCZ, where risk alleles in many genes, as well as non-coding regions of the genome, across the full allelic frequency spectrum are involved, and likely act in interacting networks (*4*). Therefore, given the large number of genes potentially involved, it has been challenging to identify which are the core set of genes that play major roles in the pathways or networks. It becomes even more challenging if the recent hypothesis of omnigenic model is true for SCZ, in which case almost all of the genes expressed in a disease-relevant cell type may confer risk to the disease through widespread network interactions with a core set of genes (*9*). Thus, it is imperative to identify disease-relevant core gene networks and possibly, the network master regulators (MRs), which are more likely to be targeted for therapeutic interventions (*10, 11*), if they exist. We define the term MR for transcription factors and genes inflicting regulatory effects on their targets. We have collected a comprehensive set of 2,198 genes curated from three sources (*12–14*) which is used in the following sections.

Transcriptional networks, as a harmonized orchestration of genomic and regulatory interactions, play a central role in mediating cellular processes through regulating gene expression. One of the most commonly used network-based modeling approaches of cellular processes are scale-free co-expression networks (*15*). However, co-expression networks and other similar networks are not comprehensive enough to fully recapitulate the entire underlying molecular interactions driving the disease phenotype (*16*). Despite the wide adoption of co-expression network analysis approaches, these methods have several limitations (*17*), such as the inability to infer or incorporate causal regulatory relationships, the difficulty to handle mammalian genome-wide networks with many genes, and the presence of high false positive rates due to indirect connections. In contrast, information-theoretic deconvolution techniques (*16*) aim to infer causal relationships between transcription factors and their downstream targets, and have been recently successfully applied in a wide range of complex diseases such as cancers (*18*) and neurodegenerative diseases (*19*).

Inferring disease-relevant transcriptional gene networks requires well-powered transcriptomic case-control datasets from disease-relevant tissues or cell types. For SCZ, although abnormal gene expression in multiple brain regions (*20*) and different types of neuronal cells (*21*) may be involved in disease pathogenesis, abnormalities in cellular and neurochemical functions of dorsolateral prefrontal cortex (DLPFC) have been demonstrated in SCZ patients (*22*). Furthermore, DLPFC controls high-level cognitive functions, many of which are disturbed in SCZ. The Common Mind Consortium (CMC) RNA-Seq data on DLPFC, which includes 307 SCZ cases and 245 controls, is currently the largest genomic data set on postmortem brain samples for neuropsychiatric disorders (*23*). The CMC study validates ^~^20% of SCZ loci having variants that can potentially contribute to the altered gene expression, where five of which involve only a single gene such as FURIN, TSNARE1, and CNTN4 (*23*). A gene co-expression analysis in the CMC study implicated a gene network relevant to SCZ, but does not address the question of whether a MR potentially orchestrates transcriptional architecture of SCZ. In the current study, using CMC RNA-Seq data, we reverse-engineered the regulatory processes mediating SCZ to identify critical MRs and to infer their role in orchestrating cellular transcriptional networks. Using another independent SCZ case-control RNA-Seq data on Cultured Neuronal cells derived from Olfactory Neuroepithelium (CNON) (*24*), we observed five MRs shared between the both data. Among the top candidate MRs identified from both datasets, we selected *TCF4*, a leading SCZ risk gene from a genome-wide association study (GWAS) (*5*), for empirical validation of predicted network activity using human induced pluripotent stem cell (hiPSC)-derived neurons. A general overview of both the computational and experimental stages of this study is provided in Fig. 1. Based on the results, we identified *TCF4* as a MR that likely contributes to SCZ susceptibility at the early stage of neurodevelopment.

**Fig. 1.**
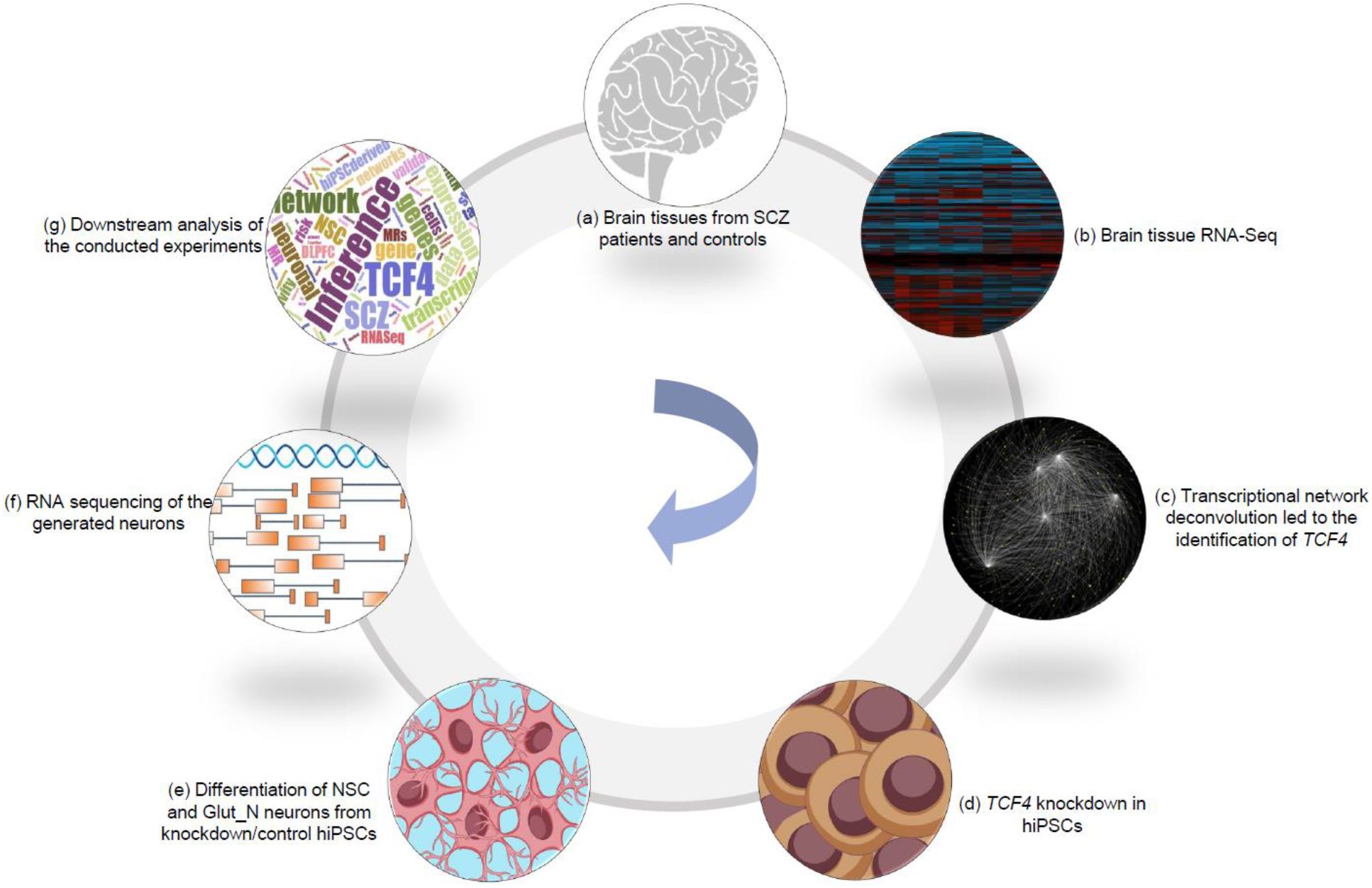
An overview of the current study. (a, b) obtaining gene expression generated from dorsolateral prefrontal cortex (CMC data) and later CNON data as an independent validation, (c) transcriptional network reconstruction based on the RNA-Seq data using ARACNe and identification of TCF4 as a disease-relevant master regulator (MR), (d) knockdown of TCF4 in human induced pluripotent stem cells (hiPSCs) derived from a patient with SCZ and two unaffected individuals, (e) differentiation of hiPSC into neural stem cells and glutamatergic neurons, (f) RNA sequencing of the generated neuronal cells with and without TCF4 knockdown, (g) downstream inferences and analysis of the conducted experiments.

## Results

### Deconvolution of transcriptional network of the CMC data uncovers master regulators

The CMC transcriptomic study implicated 693 differentially expressed genes in SCZ postmortem brains (*23*). However, the magnitude of case-control expression differences was small, posting a challenge to inferring their biological relevance. We thus reverse-engineered the transcriptional regulatory networks to infer the MRs and their transcriptional targets (regulons), from which we constructed their corresponding sub-networks. To achieve this, we employed Algorithm for Reconstruction of Accurate Cellular Networks (ARACNe) to reconstruct cellular networks (*17*) (see Materials and Methods). This approach first identifies gene-gene co-regulatory patterns by the information theoretic measure Mutual Information, followed by network pruning through removing indirect relationships in which two genes are co-regulated through one or more intermediate entities. This enabled us to observe network relationships that are more likely to represent direct transcriptional interactions (e.g., transcription factor binding to a target gene) or post-transcriptional interactions (e.g., microRNA-mediated gene inhibition). We identified 1,466 genes as hubs by computational analysis of the CMC RNA-Seq data (table S1), where the corresponding subnetworks for these hub genes contain 24,548 transcriptional interactions (table S2).

Using the reconstructed network, we next performed a virtual protein activity analysis to investigate the activity of the identified hub genes by taking into account the expression patterns of their downstream regulons through a dedicated probabilistic algorithm called VIPER (Virtual Inference of Protein-activity by Enriched Regulon analysis) (*18*). This method exploits the regulator mode of action, the confidence of the regulator-target gene interactions, and the pleiotropic features of the target regulation. We fed the generated network into VIPER to evaluate whether any of the identified hub genes in the network has significant regulatory role in the expression degrees of its downstream target genes. We then ranked the hub genes by VIPER-adjusted activity P-values (representing the statistical significance of being a MR; see Methods and Materials). We further defined a short list of genes potentially regulating a large set of targets (N=93). VIPER outputs list of genes whose activity levels is highly correlated with the expression of its regulons in the input network. However, as recommended by the VIPER developers, many of such genes are not regulators of their corresponding targets in practice. Therefore, they recommended focusing on the highly active genes in the VIPER output which are transcription factors or histone modification genes. As a result, among genes with highly significant VIPER P-values, we only took the genes from the curated list of transcription factors as potential MRs. Moreover, as suggested by the VIPER developers, we set the activity P-values at 0.005 to minimize the number of false positive MRs and to avoid obtaining candidate MRs with very few number of regulons despite high activity P-value.

To reduce false positive findings and to focus on MRs known to exert regulatory effects on their targets, we intersected the short list of potential MRs with our curated set of genes (fig. S1). This process resulted in 5 potential MRs: *TCF4, NR1H2, HDAC9, ZNF436*, and *ZNF10*. Based on previous studies (*25*), identification of the activated MRs in a disease state may be confounded, when several of its regulons are all targets of a different transcriptional factor, that is, the “shadow effect” (*25*). Because of highly pleiotropic nature of transcriptional regulations, we evaluated the shadow effects in the constructed networks. We should note that during the process of network deconvolution, most of the potential co-regulatory edges have been trimmed to preserve the scale-free topology of the network, so we expect to observe no shadow effect. As expected, no shadow connections were observed in this analysis indicating a high confidence in the constructed transcriptional network to reflect likely real regulatory connection between a MR and its targets.

### Network deconvolution of CNON data independently identifies *TCF4* as a MR

We next attempted a replication of the identified MRs in an independent RNA-Seq data set (with 23,920 transcripts) from CNON of 143 SCZ cases with 112 controls (*24*). The rationale is that if the *TCF4* network identified in CMC dataset is more of neuronal origin, confirmation of the *TCF4* MR in CNON model would add additional supporting evidence for the predicted MR given that olfactory cells are promising molecules in biological psychiatry and neuroscience (*26*). We acknowledge that that CNON culture likely contains non-neuronal cells such as epithelial cells. Because the CMC data is generated from postmortem brain tissues comprising both neuronal and non-neuronal cells, we first estimated the percentage of neuronal cells in postmortem brain DLPFC (used by CMC). Based on a recent study of cellular composition of the same brain region (DLPFC) by Lake *et al*. (*27*) in which 36,166 single cells were assayed for gene expression, we found that 26,064 cells (^~^72%) were neuronal cells, indicating that neuronal cells constitute a significant proportion of the brain cell types in postmortem DLPFC. Furthermore, out of 101 *TCF4* target genes predicted in CMC, 31 were found to have significantly higher expression in neuronal cells (enrichment P = 2.8 x 10^−10^ by Fisher’s Exact Test, fold enrichment = ^~^3.5) while the rest were all expressed in neuronal cells. We thus conclude that the constructed networks from CMC dataset stem from neuronal cells, despite possible influence from the small fraction of non-neuronal cells.

Using the similar set up criteria as used on the CMC data, for CNON data we observed 1,836 TFs or expression regulatory nodes in the constructed network, including 34,757 predicted interactions (table S3-S4). To further analyze the activity of the identified hub genes, we ran VIPER on the constructed network and identified the top MRs. The five identified MRs in the CMC network (*TCF4, NR1H2, HDAC9, ZNF436*, and *ZNF10*) were observed to be MRs within the network created from CNON data. All of these five MRs were significant in the both CMC and CNON data (FDR values are provided in table S11). Based on the deconvolution analysis, these five MRs were predicted to regulate the expression of 101, 68, 36, 66, and 43 regulons combined, respectively (table S5). Among these genes, *TCF4* is a well-established SCZ-associated risk factor identified by GWAS (*5*), which has been validated in several additional studies (*28, 29*). Rare variants in *TCF4* have also been implicated in Pitt Hopkins Syndrome and intellectual disability (*30*). However, its regulatory mechanisms at a systems level is poorly understood. Although the other identified MR may also potentially contribute to SCZ pathogenesis, given the prior biological knowledge on *TCF4*, we have focused on *TCF4* for additional analysis.

To further evaluate the connection between *TCF4* dysregulation and SCZ, we conducted pathway enrichment analysis using WebGestalt (*31*). The predicted TCF4-associated regulons were enriched for several pathways such as Notch signaling pathway (FDR= 1 x 10^−4^) and long-term depression (FDR= 8 x 10^−4^). Additionally, to check whether the promoter region of predicted *TCF4* targets are enriched for *TCF4* binding motif sequence, we conducted TF binding enrichment analysis (TFBEA) using JASPAR (*32*). The *TCF4* motif sequence is extracted from JASPAR and is used for binding enrichment analysis (Fig. 2a). We extracted the sequence of the target genes 2 kb upstream and 1 kb downstream of their Transcription Start Sites (TSS), and used a significance threshold of 0.80 to test the relative enrichment scores of its target genes. We found that the observed binding enrichment score is significantly higher than expected by chance (two-sided t-test *P*=2.2 × 10^−16^), based on a random list of genes outside of the *TCF4* subnetwork (Fig. 2b), suggesting that the predicted *TCF4* regulons can be directly targeted by *TCF4* binding events.

**Fig. 2.**
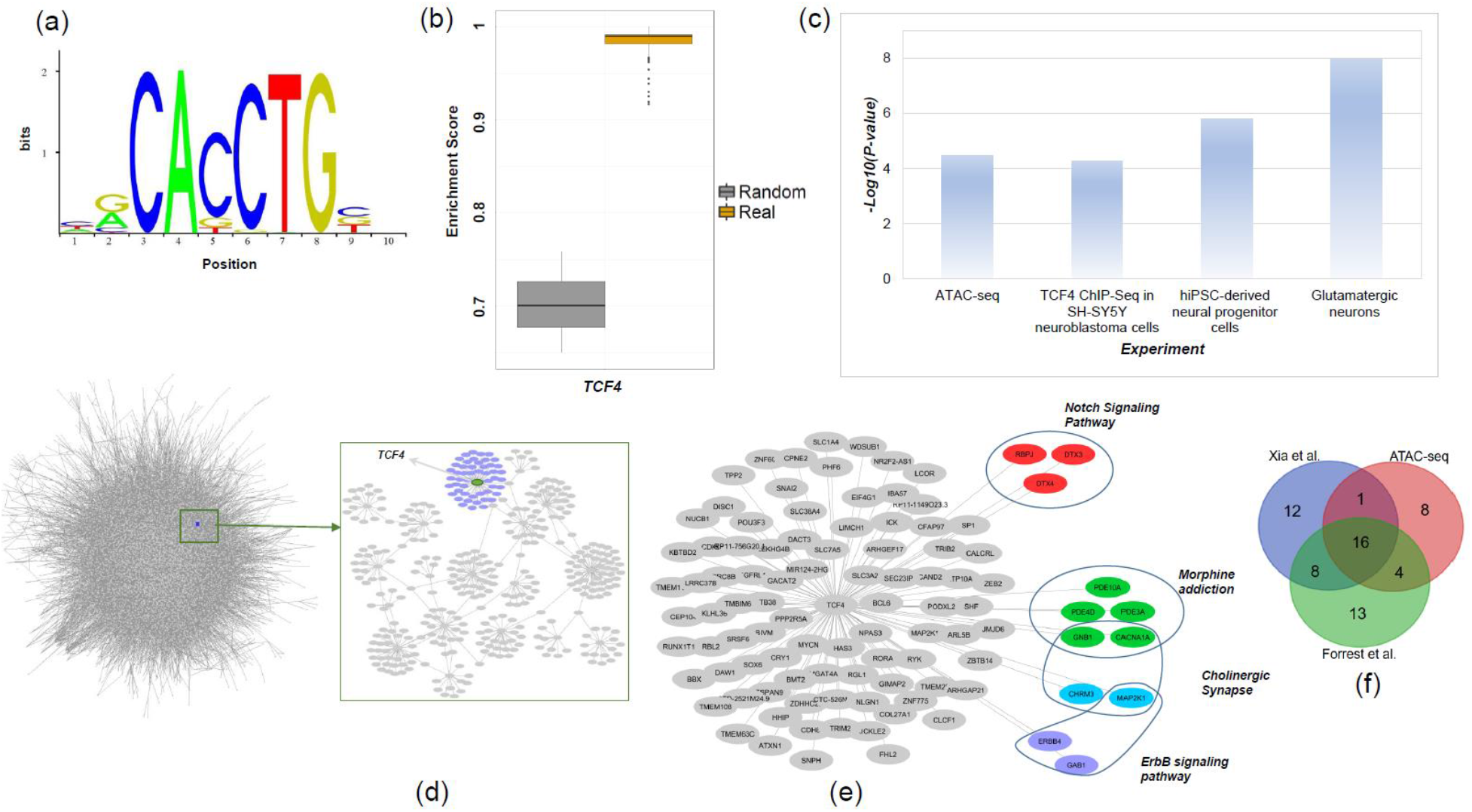
Summary of TCF4 regulons in the CMC and CNON data. (a) TCF4 motif from JASPAR database used in the analysis; (b) TCF4 binding site enrichment scores (based on TCF4 motifs from the JASPAR database) among the TCF4 regulons compared to that of a set of random genes; (c) Enrichment p-values of the TCF4 regulons, compared with several gene sets, including the predicted TCF4 sites from ATAC-Seq, the differentially expressed genes from our TCF4 knockdown experiments, and the predicted TCF4 binding sites in neuroblastoma cells by ChIP-Seq (P-value obtained from Fisher’s exact test); (d) a schematic view of the network created from CMC data as well as the TCF4 targets; (e) TCF4 targets from CMC and CNON data combined with some of the associated pathways; (f) List of overlapping predicted TCF4 targets with an ATAC-seq on hiPSC-glutamatergic neurons and two ChIP-seq datasets.

### Predicted *TCF4* regulons are significantly enriched for *TCF4* targets from ATAC-seq and ChIP-seq experiments

Our observed enrichments of TF binding motif of *TCF4* in their regulons was purely a DNA sequence-based prediction, so we next tested whether *TCF4* can physically interact with its regulons. To address this question, we first identified the target genes of *TCF4* by examining the empirical TF occupancy (i.e, TF-binding footprints) inferred by PIQ tool (*33, 34*) from our previous Assay for Transposase Accessible Chromatin (ATAC-Seq) data in human induced pluripotent stem cell (hiPSC) and hiPSC-derived glutamatergic neurons (*34, 35*). 29 target genes out of 101 regulons of *TCF4* were found to be bound by *TCF4* in ATAC-seq from Glut_Ns (total *TCF4* targets in ATAC-seq data in Glut_Ns = 3,276) (enrichment P = 3.3 x 10^−5^ by Fisher’s Exact Test, fold_enrichment = ^~^1.93), whereas only one *TCF4* target was bound by *TCF4* in hiPSCs (total *TCF4* targets in ATAC-seq data in hiPSCs = 172) (no enrichment, P = 0.417 by Fisher’s Exact Test).

Furthermore, with the list of empirically determined *TCF4* targets in a recent *TCF4* ChIP-Seq study on human SH-SY5Y neuroblastoma cells (*36*) (a total of 10,604 peaks for 5,436 unique candidate target genes), we observed an overlap of 41 *TCF4* target between *TCF4* ChIP-seq data and our inferred *TCF4* regulons, representing a significant enrichment (*P*=5.28 x 10^−5^ by Fisher’s exact test, fold change=^~^1.65). Recently, Xia *et al*. (*37*) have examined the genes regulated by *TCF4* in neural-derived SH-SY5Y cells using ChIP-seq. They have reported a total of 11,322 peaks for 6,528 unique candidate target genes. We observed 37 genes overlapping with the *TCF4* predicted target genes (*N*=101, *P*=3 x 10^−3^ by Fisher’s exact test, fold change=^~^1.5). Altogether, both ATAC-Seq and ChIP-Seq analyses demonstrated that our list of predicted regulons are significantly enriched for *TCF4* transcriptional regulatory targets, reflecting the differences in different cell types (Fig. 2c). To further probe these overlapping genes, we compiled the predicted targets of *TCF4* with the above ChIP-seq and ATAC-seq dataset on glutamatergic neurons (table S10, Fig. 2f). We observed that a large fraction of *TCF4* targets are shared between two or all of the compared datasets, which is a strong indication on the reliability of the inferred *TCF4* targets during network construction.

### Network perturbation by *TCF4* knockdown in hiPSC-derived neurons supports *TCF4* as a MR and a regulator of neurodevelopment

To further investigate the functional relevance of *TCF4* dysregulation in schizophrenia, we examined how the predicted *TCF4* transcriptional subnetwork changes with decreased *TCF4* expression in hiPSC-derived neuronal cells. We acknowledge that similar experiments have been performed on the SH-SY5Y neuroblastoma cell line (*36, 38*), which is a cell type that may be less relevant to SCZ. We first derived relatively pure NPCs (NESTIN+/PAX6+/SOX2+) from hiPSCs (TRA-1-60+/SSEA4+/NANOG+/OCT4+) (Fig. 3a-d), which were further differentiated into glutamatergic neurons (Glut_N) (VGLUT1+/MAP2+) (Fig. 3e). hiPSCs were captured from a patient with SCZ (see Methods and Materials). To capture possible differential effects of *TCF4* on gene expression during different developmental stages, we carried out *TCF4* knockdown by lentiviral shRNA in NPCs of day 3 after initiating neural differentiation (Fig. 3c-d) and in early stage of Glut_N of day 14 after initiating differentiation (Fig. 3e). We achieved significant *TCF4* knockdown in both cell stages: *TCF4* expression in the knockdown group was reduced to 32.2% and 48.6% of that in the control shRNA groups, respectively (as measured by qPCR; Fig. 3f-g, P < 0.001). We also performed RNA-Seq analysis on the same set of cells with three biological replicates at both time points in both cell types, and found that the reduction of *TCF4* transcript level was consistent with the qPCR results (Fig. 3j).

**Fig. 3.**
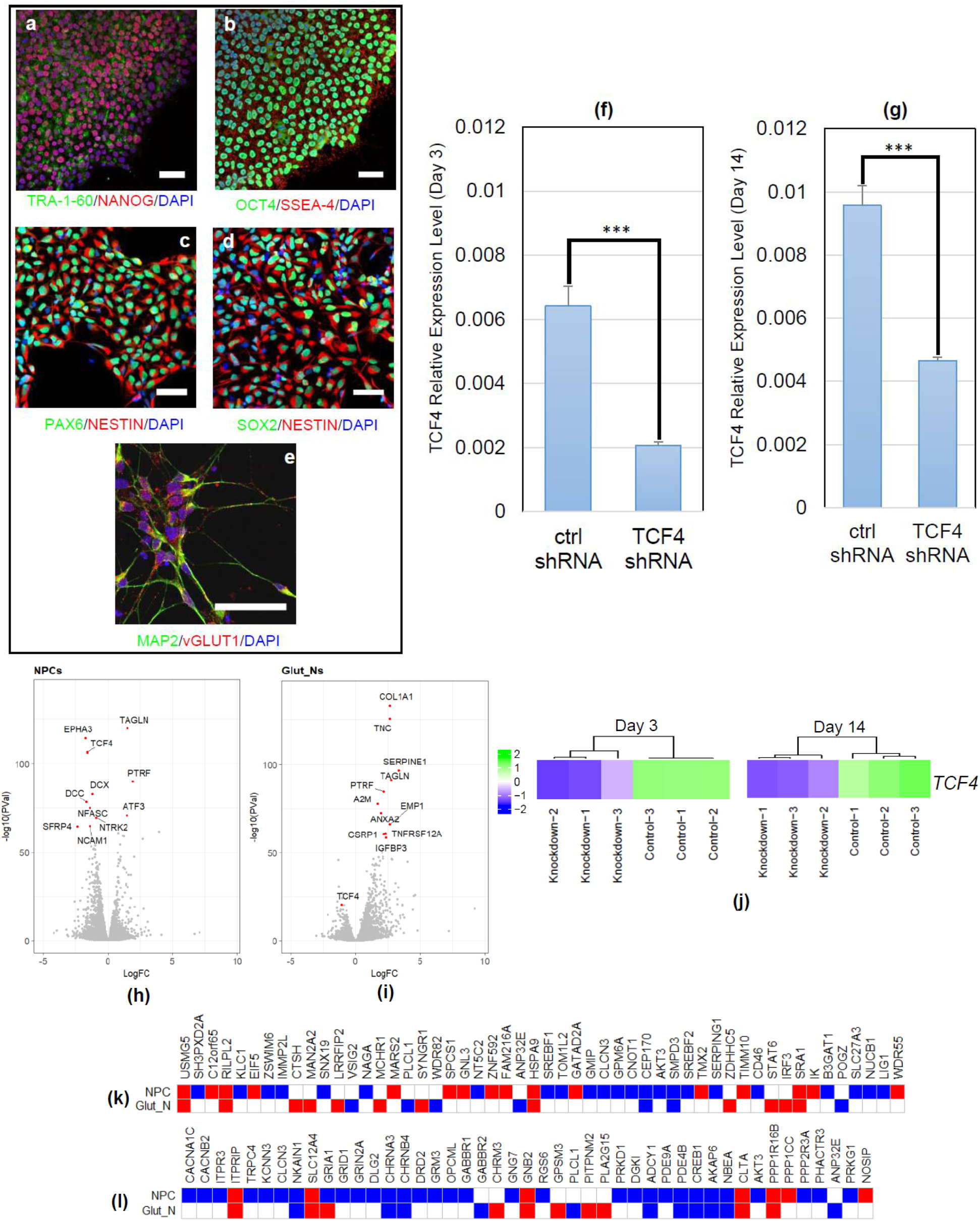
TCF4 knockdown in hiPSC-derived neuronal precursor cells (NPCs) and glutamatergic neurons (from a SCZ patient line). (a) Representative immunofluorescence (IF)-staining images of hiPSCs. hiPSCs are stained positive for pluripotency markers TRA-1-60 (green) and NANOG (62); (b) Representative immunofluorescence (IF)-staining images of hiPSCs. hiPSCs are stained positive for OCT4 (green) and SSEA-4 (62); (c) Representative IF-staining images of NPCs (day 3 post plating cells): NPCs are stained positive for PAX6 (green) and NESTIN (62); (d) Representative IF-staining images of NPCs (day 3 post plating cells): NPCs are stained positive for NESTIN (62) and SOX2 (green); (e) Representative image of day-14 neurons stained positive for MAP2 (green) and vGLUT1 (62); (f) TCF4 knockdown efficiency measured by qPCR in NPCs (3 days after plating); (g) TCF4 knockdown efficiency measured by qPCR in glutamatergic neurons (day 14 post neuronal induction); (h) volcano plot of DE genes in NPCs; (i) volcano plot of DE genes in Glut_Ns; (j) TCF4 expression levels at days 3 and 14 in RNA-Seq data (k): overlap of the DE genes upon TCF4 knockdown with a list of GWAS-implicated SCZ risk genes (up/down regulated genes are shown in red/blue, respectively. White cells indicate that the gene is not DE); (l) overlap of the DE genes upon TCF4 knockdown with a list of credible GWAS risk loci (up/down regulated genes are shown in red/blue, respectively. White cells indicates that the gene is not DE). Scale bars: 50 um in all images, cell nucleus are stained with DAPI (blue) (a-e), GAPDH is used as the endogenous control to normalize the TCF4 expression for qPCR, Error bars: mean ± SD (n=4). ***: P < 0.001, Student’s t-test, two-tailed, heteroscedastic.

Transcriptomic analysis of the RNA-Seq data of *TCF4* knockdown revealed 4,891 DE genes in the day 3 group (NPC data, Fig. 3h), including 2,330 upregulated and 2,561 downregulated genes (FDR<0.05) (see Materials and Methods). Day 14 group (Glut_N data, Fig. 3i) showed 3,152 DE genes, of which 1,862 were upregulated and 1,290 were downregulated (FDR<0.05) (table S7). To examine whether the gene network perturbed by *TCF4* knockdown in hiPSC-derived neurons is concordant with the predicted *TCF4* regulons, we compared the list of genes affected by *TCF4* knockdown and the list of 101 *TCF4* regulons from the CMC/CNON data. We found that 39 predicted *TCF4* regulons showed altered expression by *TCF4* knockdown in NPCs (enrichment P=1.55 x 10^−6^ by Fisher’s exact test, fold change=^~^1.75), while 33 predicted *TCF4* regulons were dysregulated after *TCF4* knockdown in Glut_N (enrichment P =1.05 x 10^−8^ by Fisher’s exact test, fold change=^~^2.28). This difference between the enrichment P-values of the conducted experiments compared to the neuroblastoma cell line confirmed the importance of tissue-specific investigation of transcriptional regulation (even though the set of regulons may be more conserved across tissues), and further supported the role of *TCF4* as a MR in a transcriptional network (Fig. 2d-e) relevant to neurodevelopment (Fig. 2c).

As our observed transcriptomic changes were from a SCZ patient line, we further carried out *TCF4* knockdown in cells derived from hiPSC lines of two unaffected individuals (cell lines CD07 and CD09, Fig. 4a-d) to account for variations due to individual genetic backgrounds. Because *TCF4* regulons were initially found more enriched in the list of genes perturbed by *TCF4* knockdown in glutamatergic neurons (Fig. 2c), we have focused on examining the robustness of our findings on Glut_Ns with 3 independent cultures for each condition. Using qPCR, we examined the knockdown efficiency in both cell lines (^~^40%) (Fig. 4e-f). We also confirmed the relatively high homogeneity of the differentiated neurons (80-85%) (Fig. 4g). Further transcriptomic analysis from bulk RNA-seq data also confirmed the successful reduction in *TCF4* expression in cell line CD09 (Negative Binomial Test, FDR = 1.55 x 10^−14^) as well as cell line CD07 (FDR = 2.35 x 10^−12^) (Fig. 4h). During quality assessment procedure, we removed one sequenced sample as outlier from downstream analysis (fig. S10, see Methods and Materials). In total, we identified 2,304 DE genes in CD09 and 1,744 DE genes in CD07 (FDR < 0.05). Next, we examined the enrichment of *TCF4* network regulons in the list of DE genes from the both cell lines. We observed 22 predicted *TCF4* regulons to be dysregulated after *TCF4* knockdown in Glut_N of CD09 (enrichment P = 6.85 x 10^−4^ by Fisher’s exact test, fold_enrichment = ^~^2.1) and 17 predicted *TCF4* regulons in CD07 (enrichment P = 2 x 10^−3^ by Fisher’s exact test, fold_enrichment = ^~^2). Moreover, we compared the log fold change of the genes in the two generated cell lines (CD07 and CD09) and compared them with the initial data on Glut_Ns (Fig. 4k-l). To check the consistency of cellular responses to the *TCF4* knockdown, we compared the log fold changes of the genes in CD07 and CD09 lines with the initial SCZ Glut_N data. We observed a correlation coefficient of 0.54 and 0.50 for CD07 and CD09, respectively, between the log fold changes of gene expression (P=10^−15^ and 10^−14^ for CD07 and CD09, respectively), which further supports our findings, implying that *TCF4* knockdown had a consistent effect on the transcriptome change in the several cell lines used in our study. All together, these observations re-confirmed the role of *TCF4* as MR in the created regulatory networks across individuals.

**Fig. 4.**
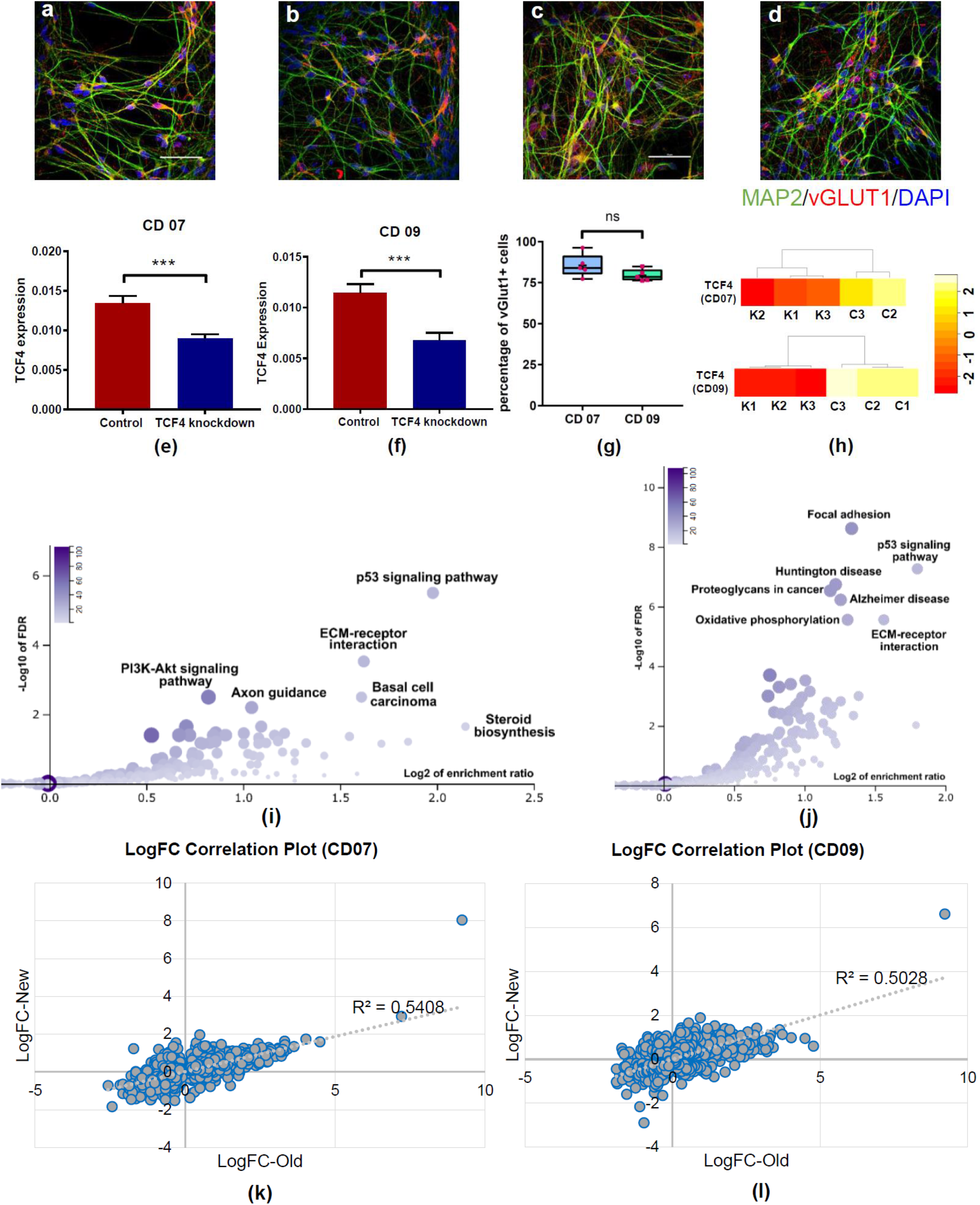
TCF4 knockdown in hiPSC-derived glutamatergic neurons in two independent cell lines CD07 and CD09 from unaffected control subjects. (a-b) IF staining of hiPSC-derived glutamatergic neurons from CD07 control line; (c-d) IF staining of hiPSC-derived glutamatergic neurons from CD09 control line; MAP2: green; vGlut1: red; DAPI: blue. Scale bar=50μM; (e-f) qRT-PCR result of TCF4 expression level in two different lines, demonstrating knock down efficiency, GAPDH is used as the endogenous control to normalize the TCF4 expression for qPCR, Error bars: mean ± SD (n=6-8) ***: P < 0.001, Student’s t-test, two-tailed, heteroscedastic; (g) Percentage of vGlut1 positive cells derived from two different hiPSC lines, cell counts were from 5 images in each line; (h) TCF4 expression levels in knockdown (K1, K2, K3) and control samples (C1, C2, C3) in the generated RNA-seq data on CD07 and CD09; (i) Pathway enrichment analysis results on the identified DE genes in CD07 (color legend indicates the number of overlapped DE genes with the corresponding pathway); (j) Pathway enrichment analysis results on the identified DE genes in CD09 (color legend indicate the number of overlapped DE genes with the corresponding pathway); (k) correlation plot of the log fold changes of the Glut_N from the SCZ cell line (“old”) and CD07; (l) correlation plot of the log fold changes of the Glut_N from the SCZ cell line (“old”) and CD09.

Although our deconvolution analysis identified 101 genes in the *TCF4* subnetwork, we reasoned that it is likely part of a much larger transcriptional network that would contain neurobiological pathways relevant to schizophrenia. Previous studies in mice (*39*) and neuroblastoma cells (*38*) have demonstrated that *TCF4* dysregulation affects a large number of genes and pathways; however, *TCF4*-related pathways have never been explored in human NPCs or glutamatergic neurons, a much stronger model for studying psychiatric disorders. To uncover the cellular pathways regulated by *TCF4* in these cell types we performed gene ontology (GO) and pathway analyses on the genes perturbed by *TCF4* knockdown.

By using WebGestalt (*31*) we identified a number of shared gene pathways, including focal adhesion, axon guidance, MAPK signaling pathway, and apoptosis (table S6 and fig. S2-S6). In table S6, we showed the altered pathways at the both time points as a result of *TCF4* knockdown, and the detailed list of up/down regulated genes in each pathway. We found almost similar enrichment for “axon guidance” (FDR =1.521 x 10^−6^ and fold enrichment=1.77 in NPCs vs FDR =5.941 x 10^−5^ and fold enrichment=1.81 in Glut_Ns), while observing some unique pathways (long-term potentiation, neurotrophin signaling, and mTOR signaling) relevant to neuronal activity. An independent pathway analysis using Metacore (see Methods and Materials, fig. S7-S8) also showed consistent results. The top gene expression changes in NPCs converged on early neurodevelopmental processes such as axon guidance, neurogenesis and attractive and repulsive receptors, which guide neuronal growth and axon targeting during development. In contrast, the most significant categories in Glut_Ns involved cell adhesion and cell-matrix interactions, suggesting that *TCF4* regulates different subsets of genes during development. In parallel, the MetaCore analysis revealed that the upregulated genes had distinct functions in NPC and Glu_Ns, involving mRNA translation and cell-matrix interactions respectively. However, down regulated genes shared a number of categories including axon guidance being the most significant pathway.

For both upregulated and downregulated genes by *TCF4* knockdown, NPCs showed more enriched neuronal-activity related gene pathways than in Glut_Ns and with greater significance (table S8-S9). It is also noteworthy that most altered neuronal-related pathways were enriched in the down-regulated set of genes upon *TCF4* knockdown. These results suggested that *TCF4* gene targets act at early stage of neurogenesis and may be more relevant to SCZ biology. We conducted a pathway enrichment analysis (Fig. 4i-j) on our independently sequenced cell lines from unaffected individuals, which yielded very similar outcomes compared to the altered pathways upon *TCF4* knockdown in the Glut_Ns in the SCZ cell line. These similarly altered pathways such as focal adhesion and p53 signaling pathway indicate that despite cross-individual genetic variations, *TCF4* plays a critical role in re-wiring the transcriptional circuitry of SCZ.

### Higher *TCF4* gene network expression activity in NPCs may be more reflective of SCZ case-control difference in prefrontal cortex

Although analyses of *TCF4* knockdown in NPC and Glut_N supported the role of *TCF4* as a MR, the *TCF4* gene network (interactome) expression activity, i.e., the correlation between *TCF4* expression and its regulon’s expression patterns, may be different in NPCs and Glut-N cellular stages. Such cell-type specific expression of the network activity may inform when the temporal expression of *TCF4* is most relevant to SCZ. Towards this end, we carried out gene set enrichment analyses (GSEA) in the CMC/CNON data, as well as the *TCF4* knockdown RNA-Seq data in NPCs and in Glut_Ns. We found that the SCZ case/control expression differences of *TCF4* regulons in CMC data (i.e., prefrontal cortex) appeared to be more correlated with the *TCF4* regulons expression changes upon *TCF4* knockdown in NPCs than in Glut_Ns. To further demonstrate the closer similarity of the *TCF4* interactome expression activity in prefrontal cortex and in NPCs, we depicted the GSEA enrichment scores in three data sets (CMC/CNON, NPC and Glut_N) in a Circos plot (fig. S9a). We found that the pattern of SCZ case/control expression differences of *TCF4* and its regulon in prefrontal cortex appeared to be more similar to the pattern of higher *TCF4* network expression activity in NPCs than in Glut_Ns. Similarly, the GSEA rank metric score of the *TCF4* regulons also showed that the SCZ case/control differential *TCF4* interactome expression pattern in prefrontal cortex was more comparable to that in NPCs than in Glut_Ns (fig. S9b). We acknowledge that prefrontal cortex, from which CMC data is generated, largely consists of differentiated cells. In this regard, our observation on higher *TCF4* expression and its positively correlated regulon expression activity in NPCs is somewhat unexpected. One explanation could be that *TCF4* expression peaks during early cortical development (*40*). One of the limitations of our observation is that both Glut_N and NPC stages represented early neurodevelopmental stage, which would not recapitulate the actual gene network in adult brain prefrontal cortex. Nonetheless, our observation is intriguing and may arguably imply that *TCF4* may be a SCZ risk factor at early stage of neurodevelopment and neurogenesis, which warrants further investigation.

### *TCF4* gene network in NPCs may be more relevant to SCZ disease biology

Given that the expression pattern of *TCF4* and it regulons in prefrontal cortex are more correlated to NPCs than Glut_Ns (fig. S9a), we reasoned that *TCF4* gene network in NPCs may be more relevant to SCZ disease biology. To test this hypothesis, we have examined the enrichments of genes regulated by *TCF4* in NPCs, as compared to Glut_Ns for disease-relevant gene pathways, SCZ GWAS risk genes, and *de novo* SCZ mutations. We conducted a series of comparative tests. To test whether the *TCF4* gene network acting in NPCs is more relevant to the genetic etiology of SCZ, we compared a list of known SCZ susceptibility genes from genome-wide significant risk loci (*5*) that showed gene expression change upon *TCF4* knockdown in NPCs and in Glut_Ns. We found that 52 of those genes (out of 148 reported in reference (*5*)) were altered by *TCF4* across at least one of two time points in either NPCs or Glut_Ns. Of these 52 SCZ credible genes that showed TCF4-altered expression, 63.5% were uniquely found in NPCs (*N*=33, Fisher’s Exact test= 0.019, enrichment factor=1.5) while 25% (*N*=13, Fisher’s Exact test= 0.950, enrichment factor=0.6) were uniquely found in Glut_Ns (Fig. 3k). Furthermore, for the 92 SCZ-associated single nucleotide variants (SNVs) that were indexed for genes related to neuronal electrical excitability and neurotransmission (*41*), 45 had altered expression by *TCF4* knockdown either in NPCs or Glut_Ns. Among them, 24 genes (53.34%, Fisher’s Exact test: 0.021, enrichment factor=1.17) were uniquely found in NPCs while only 7 genes (15.6%, Fisher’s Exact test: 0.90, enrichment factor=0.54) were uniquely found in Glut_Ns (Fig. 3l). Similarly, we compiled 176 candidate SCZ risk genes from the CLOZUK GWAS study (*42*), and 54 genes were altered by *TCF4* in either NPCs or Glut_Ns where 72% (*N*=39, Fisher’s Exact test: 0.035, enrichment factor=1.1) of the genes were uniquely found in NPCs compared to ^~^17% (*N*=9, Fisher’s Exact test: 0.98, enrichment factor=0.36) of the genes uniquely found in Glut_Ns. Lastly, we conducted SCZ *de novo* SNV (dnSNV) enrichment analysis to probe the overlap of TCF4-associated gene expression changes and dnSNV previously reported in SCZ (Fig. 5a) using a list of *de novo* SNVs from SCZ patients and control subjects (see Methods and Materials). We found that while TCF4-affected genes in NPCs showed significant enrichment for both loss-of-function (LoF) and missense SCZ dnSNVs, and coding dnSNVs in general (Fig. 5b,d-, only LoF dnSNVs were enriched in Glut_Ns (Fig. 5c-d). The analyses of both common variants and dnSNVs associated with SCZ in TCF4-affected genes nominated NPC as a more disease-relevant cell-type and stage of development for *TCF4* gene network. Altogether, our results suggested that *TCF4* and its regulons in early stage of neurodevelopment and neurogenesis may be more relevant for SCZ pathogenesis. However, additional studies should be conducted to further validate this hypothesis.

**Fig. 5.**
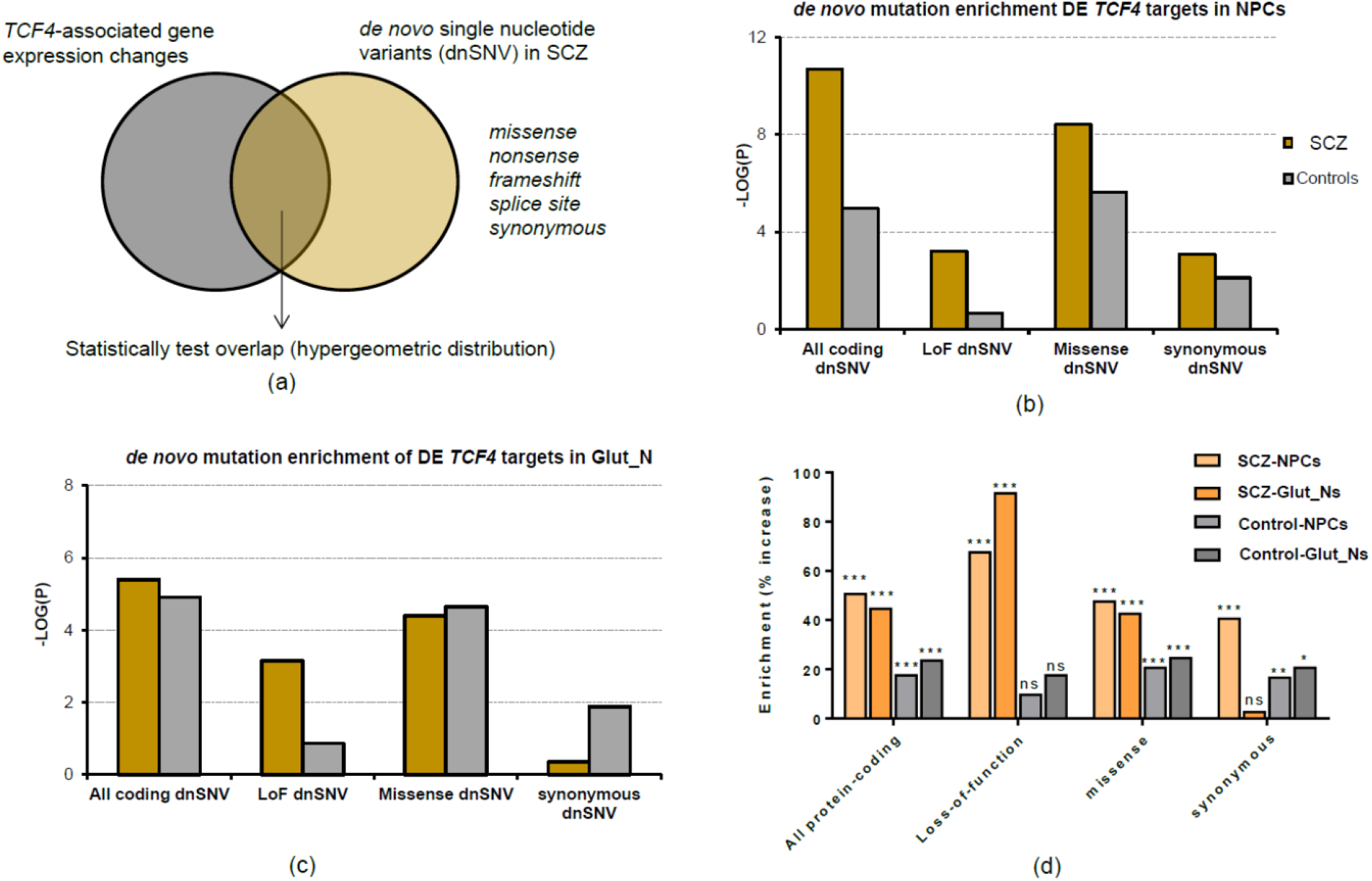
Enrichment of SCZ de novo SNVs in TCF4-associated genes expression changes in SCZ samples and control subjects. (a) analysis framework; (b) Enrichment P-values (−log_10_ transformed) of all DE genes at day 3 (NPCs), The P-value on the y-axis was obtained from a hypergeometric test was used to test the statistical significance of each overlap (number of shared genes between lists) and using all protein coding genes as a background set; (c) Enrichment P-values (−log_10_ transformed) of all DE genes at day 14 (Glut_N); (d) Fold change values of SCZ dnSNVs in TCF4-associated gene expression changes.

## Discussion

Like other complex disorders, SCZ is polygenic or even possibly omnigenic (*9*) with hundreds or even thousands of susceptibility genes each with small effect size. A major challenge for understanding the biological implications of genetic findings is to identify the core gene networks in disease relevant tissues or cell types (*9*). In this study, with the aim of uncovering the gene expression-mediating drivers that may contribute to SCZ pathogenesis, we have employed an approach to assemble tissue-specific transcriptional regulatory networks on two independent RNA-Seq datasets from dorsolateral prefrontal cortex (CMC data) and cultured cells derived from olfactory neuroepithelium (CNON data). CNON was used as a validation data sets, to evaluate whether findings from brain samples can be seen from *in vitro* cell culture that contains neuronal cells. We identified *TCF4* as a MR based on the deconvolved transcriptional networks from the two independent data sets. For the predicted gene network of *TCF4*, a known leading SCZ GWAS locus, we have empirically validated the predicted *TCF4* gene targets by analyzing both TF-binding footprints in neuronal cells and by transcriptomic profiling of TCF4-associated gene expression changes in both hiPSC-derived NPCs and Glut_Ns upon *TCF4* knockdown.

Compared to the commonly used weighted gene co-expression network analysis (WGCNA) (*43*), the method used in our study aims to find MRs rather than identifying co-expression patterns as well as bearing a number of unique features, including the ability to elucidate potentially causal interactions, to illustrate the hierarchical structure of the identified transcriptional networks, and to identify potential feed-forward loops between MRs that may cause synergistic effects in the network. The confidence of the identified MRs and their subnetworks was further strengthened by their presence in both transcriptomic datasets, and experimentally validated in hiPSC-derived neuronal cells using ATAC-Seq data, ChIP-Seq data, and transcriptomic profiling.

*TCF4* is one of the leading SCZ risk genes identified by GWAS (*5*). However, the molecular mechanism underlying the genetic association remain elusive. Here, we have shown computationally that *TCF4* is one of the top MRs of SCZ, and this was empirically validated in hiPSC model of mental disorders. Although the downstream gene targets of *TCF4* have been previously profiled in a neuroblastoma cell lines (*38*), the hiPSC cellular model used here enabled us to examine the neurodevelopmental relevance of *TCF4* gene network. By examining the TCF4-pertubred gene networks in hiPSC-derived NPC (early neuronal development/neurogenesis stage) and in differentiated Glut_N, we found that TCF4-altered genes in NPC are more enriched for gene pathways that are related to neuronal activities. Additional experiments on two independent cell lines from unaffected individuals further validated our findings as to how *TCF4* targets a large body of genes in Glut_Ns. Furthermore, we have shown that TCF4-altered genes in NPC are more enriched for credible SCZ GWAS risk genes and for SCZ-associated dnSNVs. Thus, multiple lines of evidence from our study suggest that *TCF4* gene network expression activity in the early stage of neurodevelopment and neurogenesis may be more important for SCZ pathogenesis. In an attempt to further substantiate our findings on elucidating the transcriptional effects of *TCF4*, we combined the data from all three hiPSC lines (1 SCZ case and 2 controls lines) and identified differentially expressed genes upon *TCF4* knockdown in Glut_Ns. This allowed us to boost the power of our analysis where we observed 5,584 DE genes (FDR<0.05) with 3017 upregulated and 2567 downregulated genes. Notably, we observed 47 DE genes overlapping with the 101 predicted *TCF4* regulons (Fisher’s Exact test: p=1.3 x 10^−5^, fold_enrichment= 1.83). This analysis further demonstrates the significance of our findings on the regulatory role of *TCF4* on its regulons.

Although our empirical validation of MR has focused on *TCF4*, other MRs that we identified here suggest that additional gene networks are relevant to SCZ. Indeed, the identified MRs contain many of the TFs previously reported in the literature that may be potentially involved in the pathogenesis of SCZ, such as *JUND* (*44*), *NRG1* and *GSK3B* (*45*), *NFE2* and *MZF1* (*46*). Among those newly identified MRs, methylation patterns of *TMEM9* (*47*) has been reported to be associated with Parkinson’s disease, yet *CRH* is a protein-coding gene associated with Alzheimer’s disease, depression, and SCZ (*48*), and *ARPP19* has been identified to be associated with nerve terminal function (*49*). A particularly noteworthy new candidate MR is *HDAC9*, a histone deacetylase inhibitor that is important for chromatin remodeling. It has been shown to regulate mouse cortical neuronal dendritic complexities (*50*) as well as hippocampal-dependent memory and synaptic plasticity under different neuropsychiatric conditions (*51*). In our deconvoluted gene network of *HDAC9*, its regulons were moderately enriched for genes related to focal adhesion (*P*=0.0115), a biological function that has been shown to be altered in SCZ (*52*). Of note, *TCF4*-altered genes in both NPCs and Glut-Ns also showed enrichment of focal adhesion pathway. The MRs we identified, and their subnetworks may provide a rich resource of SCZ-relevant gene networks that can be perturbed to increase our understanding of SCZ disease pathophysiology. Moreover, we showed that despite cross-individual genetic variations that may drive different expression patterns, *TCF4* appears to be a strong SCZ MR in three different cell lines.

The selection of *TCF4* as top MR candidate for validation was based on the consistency between CMC data and CNON data sets. However, we noted that *TCF4* regulons predicted by CMC data and CNON data were different. This difference may have largely arisen from the distinct cellular composition of the prefrontal cortex in CMC data and the primary olfactory neuroepithelium in CNON data. CMC data was generated from post-mortem human prefrontal cortex in which tens of different cell-types might have been involved. On the other hand, CNON data which was used as a complementary data aimed at further probing the in-silico effects of MRs on their targets, is less heterogeneous but with different cellular composition from the prefrontal cortex in CMC. We acknowledge that CNON could be a mixture of dividing neuronal cells and even epithelia cells. Nonetheless, our transcriptomic comparative analysis (fig. S11) indicated that transcriptomic profiles of CNON, CMC samples and our iPSC-derived NPCs were all moderately to highly correlated with each other and with GTEx brain frontal cortex and hippocampus (*53*), and CNON resembles most to CMC data compared to NPCs. Alternatively, the different *TCF4* regulons from the two different datasets may be due to the stochastic nature of network deconvolution approaches in general when sample size is moderate. The original developers recommended >100 samples in any analysis to reduce variation, and we had more than this recommended number in both data sets. Finally, it is noteworthy that previous work also observed similar differences of predicted regulons when analyzing different cancer data sets from tissues and from cell lines (*54*). Thus, future network deconvolution in single cells within each cell type may minimize such variations.

Despite the possible effects of different cellular compositions on *TCF4* regulons as discussed above, we have focused on our empirical validation in hiPSC-derived glutamatergic neurons. This is largely based on recent findings that show glutamatergic neurons to be one of the most relevant cell types to SCZ (*55, 56*). However, other cell types such as interneurons and microglia have also been implicated in pathogenesis of SCZ. Given cell type-specific expression profiles, the transcriptomic effects of *TCF4* knockdown in different cell types are likely different to some extent, which would be an interesting future research subject. Furthermore, although our relatively homogenous 2D neuronal cultures (80-95%) have their advantages, they may not reflect the *in vivo* brain circuit where different cell types interact with each other. Therefore, it would be interesting to interrogate possible cell type-specific effects of *TCF4* knockdown in a mixed cell culture or using brain organoids, followed by single-cell RNA-seq (scRNA-seq) analysis.

In summary, by employing both a computational approach and a hiPSC neuronal developmental model, we have identified *TCF4* as a MR that likely contributes to SCZ susceptibility at the early stage of neurodevelopment. Although powerful, we acknowledge the limitation of our approach in identifying top MRs, because selecting a top MR is not purely “data-driven”; for instance, *TCF4* was identified as a MR among many other MRs and our selection of *TCF4* as the top MR for validation was not only “data-driven” but also based on prior knowledge about the *TCF4* association with SCZ. Nonetheless, our study suggests that MRs in SCZ can be identified by transcriptional network deconvolution, and that MRs such as *TCF4* as well as other MRs may constitute convergent gene networks that confer disease risk in a spatial and temporal manner. Therefore, *TCF4* and other identified MRs may be a building block of SCZ polygenic/omnigenic architecture which collectively drive SCZ onset and progression. Transcriptional network deconvolution in larger and more SCZ-relevant cell types/stages, combined with empirical network perturbation, will further deepen our understanding of the genetic contribution to SCZ disease biology.

## Methods and Materials

### Pre-processing of the RNA-Seq data sets

RNA-seq data of DLPFC (Dorsolateral Prefrontal Cortex) Release 1.2 was downloaded from “normalized SVA corrected” directory of the CommonMind Consortium (CMC) Knowledge Portal (https://www.synapse.org/#!Synapse:syn5609499) using Synapse Python client (http://docs.synapse.org/python/). The data being used in this study contains 307 SCZ cases and 245 controls where the paired-end RNA-Seq data has been generated on an Illumina HiSeq 2500. Brain tissues in this dataset were obtained from several brain bank collections including: Mount Sinai NIH Brain and Tissue Repository, University of Pittsburgh NeuroBioBAnk and Brain and Tissue repositories, University of Pennsylvania Alzheimer’s Disease Core Center, and the NIMH Human Brain Collection Core. Detailed procedure of tissue collection, sample preparation, data generation and processing can be found in the CommonMind Consortium Knowledge Portal Wiki page (https://www.synapse.org/#!Synapse:syn2759792/wiki/69613).

The normalized RNA-Seq read counts from 143 SCZ cases and 112 controls extracted from cultured primary neuronal cells derived from neuroepithelium were generated as previously described (*24*). Similar to the CMC data, the raw CNON data consists of 100 bp paired-end reads. The data has been pre-processed to exclude rRNA and mitochondrial genes. Later, normalized counts were calculated by the DESeq2 software.

### Transcriptional network reconstruction

ARACNe (Algorithm for the Reconstruction of Accurate Cellular Networks), an information-theoretic algorithm for inferring transcriptional interactions, was used to identify candidate transcriptional regulators of the transcripts annotated to genes both in CMC and CNON data. First, mutual interaction between a candidate TF(*x*) and its potential target (*y*) was computed by pairwise mutual information, MI(*x,y*), using a Gaussian kernel estimator. A threshold was applied on MI based on the null-hypothesis of statistical independence (P<0.05, Bonferroni corrected for the number of tested pairs). Second, the constructed network was trimmed by removing indirect interactions the data processing inequality (DPI), a property of the MI. Therefore, for each (*x,y*) pair, a path through another TF(*z*) was considered and every path pertaining the following constraint were removed (*MI*(*x,y*) < min(*MI*(*x,z*),*MI*(*z,y*))). P-value threshold of 1 x 10^−8^ using DPI=0.1 (as recommended(*17*)) were used when running ARACNe.

### Virtual protein activity analysis

The regulon enrichment on gene expression signatures was tested by the VIPER (Virtual Inference of Protein-activity by Enriched Regulon analysis) algorithm. First, the gene expression signature is obtained by comparing two groups of samples representing distinctive phenotypes or treatments. In the next step, regulon enrichment on the gene expression signature can be computed using Analytic rank-based enrichment analysis (*56*). At the end, significance values (P-value and normalized enrichment score) are computed by comparing each regulon enrichment score to a null model generated by randomly and uniformly permuting the samples 1,000 times. The output of VIPER is a list of highly active MRs as well as their activity scores and their enrichment P-values. Further information about VIPER can be accessed in reference (*18*).

### Transcription factor binding site enrichment and footprint analysis

Human reference genome (hg19) was used to extract the DNA sequence around transcript start sites (TSSs) for transcription binding enrichment analysis. We obtained the gene coordinates from Ensembl BioMart tool (*57*) and scanned 2000 bp upstream and 1000 bp downstream of the TSS. The motifs of the TFs were obtained from JASPAR and the extracted sequences of each target were then fed into JASPAR and analyzed versus their corresponding TFs. Then, using the PWM of the TF, JASPAR employs the modified Needleman-Wunsch algorithm to align the motif sequence with the target sequence in that the input sequence is scanned to check whether or not the motif is enriched. The output is the enrichment score of the input TF in the designated target genes.

We used PIQ to assess the local TF occupancy footprint from ATAC-Seq data (*33, 34*). We extracted the corresponding BED files for TF footprint analysis using the PIQ R package. All the footprints were annotated using the TF matrix with the names of different TFs annotated in the BED files. For each sample, footprints were generated using three different PIQ purity scores (0.7, 0.8, or 0.9; equivalent to FDR of 0.3, 0.2, or 0.1, respectively). The corresponding files are then extracted using the MR list and the peak names/coordinates containing *TCF4* gene are collected as a subset of the original BED file. Such subsets of genomic coordination are then annotated using the findPeaks.pl included in the HOMER package with the hg19 reference genome.

### hiPSC generation and cell culture

The hiPSC lines used for deriving neurons were generated by Sendai virus method at Rutgers University Cell and DNA Repository (RUCDR)-NIMH Stem Cell Center from cryopreserved lymphocytes (CPLs) of the MGS collection (*58*). The patient line was derived from a 29 year-old male SCZ patient (cell line ID: 07C65853), and the other two lines were derived from unaffected controls (cell line IDs: 05C39664 and 05C43758, designated as CD07 and CD09, respectively). All three lines do not have large CNVs associated with SCZ. The NorthShore University HealthSystem Institutional Review Board (IRB) approved the study. hiPSCs were cultured using the feeder-free method in Geltrex (Thermofisher)-coated plates in mTeSR1 medium (StemCell). The media were changed daily, and cells were passaged as clumps every 5 days using ReLeSR (StemCell) in the presence of 5 μm ROCK inhibitor (R&D Systems). hiPSCs were characterized by positive IF staining of pluripotent stem cell markers OCT4, TRA-1-60, NANOG, and SSEA4 (Fig. 3a-b).

### hiPSC-derived neural progenitor cells (NPCs)

NPCs were differentiated from hiPSC using the PSC Neural Induction Medium (Thermofisher) with modifications. In brief, hiPSCs were re-plated as clumps in Geltrex (Thermofisher)-coated 6-well plates (Thermofisher) in mTeSR1 (StemCell) on Day 0. From Day 2, the medium was switched to PSC Neural Induction Medium and changed daily. On Day 11, NPCs were harvested using Neural Rosette Selection Reagent (StemCell). NPCs were maintained in Neural Expansion Medium (Thermofisher) and passaged every 4-6 day in the presence of 5 μM ROCK inhibitor (R&D Systems) until the 4^th^ passage (P4). P4 NPCs were characterized by positive IF staining for Nestin and Pax6 (Fig. 3c-d).

### hiPSC differentiation and shRNA-mediated *TCF4* (ITF-2) knockdown

hiPSC-derived NPCs were plated at the density of 3 x 10^5^ per well in a 12-well plate pre-coated with Geltrex for 2 hours. The differentiation media used for induced differentiation of NPCs contains neural basal medium, B-27 supplement, L-glutamine, 10 ng/ml BDNF, 10 ng/ml GDNF, and 10 ng/ml NT-3(*34*). NPCs were allowed to grow and differentiate for 3 or 14 days post induction, respectively. At the corresponding day, 20 μl (1.0 x 10e5 infection units of virus, IFU) of either *TCF4* or control shRNA lentiviral particles (Santa Cruz sc-61657-V for *TCF4*, which contains a pool of 3 target-specific, propriety constructs targeting human *TCF4* gene locus at 18q21.2; and sc-108080 for control, which contains a set of scrambled non-specific shRNA sequence) were added into each well (3 replicates for each group), and the transduced cells were further cultured for 48 hours before collection. Total RNAs were extracted from homogenized cell lysate using RNEasy Plus mini kit (Qiagen), and the quality of extracted total RNA were examined using Nanodrop spectrum analysis and agarose gel electrophoresis. The day-14 excitatory neurons were characterized by positive IF staining for VGLUT1 (Fig. 3e).

### qPCR validation of *TCF4* shRNA knockdown efficiency

We used qPCR to verify the knockdown efficiency of shRNA before applying RNA-Seq. Briefly, cDNAs were reverse-transcribed from total RNAs using the High-Capacity Reverse Transcription Kit with RNase Inhibitor (ThermoFisher). 50 ng of total RNA were used for each reverse-transcription reaction (20 μl) according to the manufacturer’s instructions. cDNA was diluted 15-fold before applying to qPCR. Subsequent qPCR analysis was performed on a Roche LightCycler 480 II real-time PCR machine using TaqMan probes against *TCF4* and *GAPDH* (as internal reference) respectively. Briefly, 5 μl diluted RT product was applied to each 10 μl qPCR reaction using the TaqMan Universal Amplification kit (ThermoFisher/AppliedBiosystems) with customized TaqMan probes (Hs.PT.58.21450367 for *TCF4*, Integrated DNA Technologies; and Hs03929097_g1 for GAPDH, AppliedBiosystems). Cycle parameters: 95°C, 10 min; 95°C, 15 sec, 60°C 1 min, 45 cycles. Data analysis was performed using the build-in analysis software of Roche 480 II with relative quantification (ΔCt method applied).

### RNA-Seq analysis on hiPSC-derived NPCs and Glut_Ns

RNA-seq libraries were prepared from total RNAs from the collected neuronal cells of 12 samples (three biological replicates, i.e., independent cell cultures, of four distinct groups). These sample groups are for *TCF4* expression knockdown and control at hiPSC-derived NPC stage (day 3 post-differentiation) and Glut-N stage (day 14 post-differentiation). Total RNAs were extracted as described above. Three main methods for quality control (QC) of RNA samples were conducted including: (1) preliminary quantification; (2) testing RNA degradation and potential contamination (Agarose Gel Electrophoresis); (*10*) checking RNA integrity and quantification (Agilent 2100). After the QC procedures, library was constructed and library QC was conducted consisting three steps including: (1) testing the library concentration (Qubit 2.0); (2) testing the insert size (Agilent 2100); (*10*) precise quantification of library effective concentration (qPCR). The quantified libraries were then sequenced using Illumina HiSeq/MiSeq sequencers with 150 bp paired-end reads, after pooling according to its effective concentration and expected data volume.

The paired-end sequenced reads for each sample were obtained as FASTQ files. Initial quality check of the raw data was performed using FastQC. All FASTQ files passed the quality control stage including removing low quality bases, adapters, short sequences, and checking for rRNA contamination. The average number of pre-processed reads per sample was ^~^22,740,000. High quality reads were mapped to the reference genome GRCh38 using STAR (*59*). Sorting and counting was also performed using STAR based on the procedure recommended by the developers. To detect DE genes, the output read counts were fed to edgeR software package (*60*). Normalization, batch correction, and differential expression analysis was conducted using the recommended settings.

### Pathway enrichment and GO analysis

Pathway enrichment and GO analysis were conducted using WebGestalt (*31*) v. 2017. KEGG was used as the functional database the list of expressed genes were used as the background. The maximum and minimum number of genes for each category were set to 2000 and 5, respectively based on the default setting. Bonferroni-Hochberg (BH) multiple test adjustment was applied to the enrichment output. FDR significance threshold was set to 0.05.

### MetaCore pathway analysis

Ensembl transcript IDs of differentially expressed genes were imported in Metacore (v 6.32) and processed using the ‘Enrichment Analysis’ workflow. For each time point (3 day or 14 day), all the upregulated (FDR<0.05), all downregulated (FDR<0.05) and the top 2000 differentially expressed genes (FDR<0.05) were analyzed. Each list was compared to a defined background set of genes consisting of all the expressed transcripts (FPKM>1) at the relevant time point (3 day or 14 day). The ‘Pathway Maps’ or ‘Process networks’ that were enriched in each analysis were considered significant when FDR<0.05.

### Enrichment analysis on *de novo* mutations in schizophrenia

Data on *de novo* variants in schizophrenia and controls was obtained from http://denovo-db.gs.washington.edu/denovo-db [accessed 11/04/2017] (*61*). Variants identified in schizophrenia and control subjects were stratified into four categories (all protein coding, missense, loss-of-function and synonymous) based on annotations from the denovo-db. To generate a list of variants affecting protein coding (‘all protein coding’), *de novo* variants outside exonic coding regions were excluded (including *de novos* in 5’-UTRs, 3’-UTRs, intronic and intergenic regions and non-coding exons) as well as synonymous variants. The ‘loss-of-function’ list included frameshift, stop gained, stop lost, start lost, splice acceptor and donor variants. We then compiled a total of six lists containing different sets of differentially expressed genes from the *TCF4* knockdown (KD) experiments at day 3 and day 14, as follows: day3_all_sig (all differentially expressed genes FDR<0.05; 4898 genes), day3_up_sig (all upregulated genes FDR<0.05; 2332 genes), day3_down_sig (all downregulated genes FDR<0.05; 2566), day14_all_sig (all differentially expressed genes FDR<0.05; 3156 genes), day14_up_sig (all upregulated genes FDR<0.05; 1864 genes), day14_down_sig (all downregulated genes FDR<0.05; 1292). Each of the *de novo* lists for schizophrenia and controls were intersected with the relevant list of differentially expressed genes from the *TCF4* KD experiments to determine the shared number of genes in each comparison. A hypergeometric test was used to test the statistical significance of each overlap (number of shared genes between lists) and using all protein coding genes as a background set (20,338, human genome build GRCh38.p10). The data is displayed as the −log10 (P-value) for each comparison.

## Acknowledgements

We thank the CommonMind Consortium to generate the RNA-seq data on dorsolateral prefrontal cortex from patients and control subjects (The data were generated as part of the CommonMind Consortium supported by funding from Takeda Pharmaceuticals Company Limited, F. Hoffman-La Roche Ltd and NIH grants R01MH085542, R01MH093725, P50MH080405, R01MH097276, R01MH075916, P50MH096891, P50MH084053S1, R37MH057881 and R37MH057881S1, HHSN271201300031C, AG02219, AG05138 and MH06692). We thank the developers of the ARACNe software for comments on the software usage, and thank the Wang lab members for helpful suggestions and discussions. This study was supported by NIH grant MH108728 (K.W.), HG006465 (K.W.), CHOP Research Institute (K.W.), MH086874 (O.E. and J.A.K.), MH102685, MH106575, and MH116281 (J.D.).

## Author Contributions

A.D.T. conceived the study, designed the study, performed the computational experiments, analyzed the data, and wrote the manuscript. C.A., O.V.E., T.S., and J.A.K. generated and pre-processed the CNON data and edited the manuscript. S.Z. and H.Z. generated the hiPSC cells, conducted *TCF4* knockdown and other experiments and wrote the relevant Methods sections. M.P.F. advised on the study and performed additional enrichment analysis on *TCF4*. K.W. conceived the study, wrote and edited the manuscript, and supervised the project. J.D. supervised the experimental sections of the study, wrote and edited the manuscript.

## Data Availability

All of the data generated in this study are deposited in Gene Expression Omnibus (GEO) under accession number GSE128333.

## References

1. J. McGrath, S. Saha, D. Chant, J. Welham, Schizophrenia: a concise overview of incidence, prevalence, and mortality. Epidemiol Rev 30, 67–76 (2008).

2. R. C. Kessler, H. Birnbaum, O. Demler, I. R. Falloon, E. Gagnon, M. Guyer, M. J. Howes, K. S. Kendler, L. Shi, E. Walters, E. Q. Wu, The prevalence and correlates of nonaffective psychosis in the National Comorbidity Survey Replication (NCS-R). Biol Psychiatry 58, 668–676 (2005).

3. R. Hilker, D. Helenius, B. Fagerlund, A. Skytthe, K. Christensen, T. M. Werge, M. Nordentoft, B. Glenthoj, Heritability of Schizophrenia and Schizophrenia Spectrum Based on the Nationwide Danish Twin Register. Biol Psychiatry 83, 492–498 (2018).

4. E. Rees, M. C. O’Donovan, M. J. Owen, Genetics of schizophrenia. Current Opinion in Behavioral Sciences 2, 8–14 (2015).

5. C. Schizophrenia Working Group of the Psychiatric Genomics, Biological insights from 108 schizophrenia-associated genetic loci. Nature 511, 421–427 (2014).

6. C. R. Marshall, D. P. Howrigan, D. Merico, B. Thiruvahindrapuram, W. Wu, D. S. Greer, D. Antaki, A. Shetty, P. A. Holmans, D. Pinto, M. Gujral, W. M. Brandler, D. Malhotra, Z. Wang, K. V. F. Fajarado, M. S. Maile, S. Ripke, I. Agartz, M. Albus, M. Alexander, F. Amin, J. Atkins, S. A. Bacanu, R. A. Belliveau, Jr., S. E. Bergen, M. Bertalan, E. Bevilacqua, T. B. Bigdeli, D. W. Black, R. Bruggeman, N. G. Buccola, R. L. Buckner, B. Bulik-Sullivan, W. Byerley, W. Cahn, G. Cai, M. J. Cairns, D. Campion, R. M. Cantor, V. J. Carr, N. Carrera, S. V. Catts, K. D. Chambert, W. Cheng, C. R. Cloninger, D. Cohen, P. Cormican, N. Craddock, B. Crespo-Facorro, J. J. Crowley, D. Curtis, M. Davidson, K. L. Davis, F. Degenhardt, J. Del Favero, L. E. DeLisi, D. Dikeos, T. Dinan, S. Djurovic, G. Donohoe, E. Drapeau, J. Duan, F. Dudbridge, P. Eichhammer, J. Eriksson, V. Escott-Price, L. Essioux, A. H. Fanous, K. H. Farh, M. S. Farrell, J. Frank, L. Franke, R. Freedman, N. B. Freimer, J. I. Friedman, A. J. Forstner, M. Fromer, G. Genovese, L. Georgieva, E. S. Gershon, I. Giegling, P. Giusti-Rodriguez, S. Godard, J. I. Goldstein, J. Gratten, L. de Haan, M. L. Hamshere, M. Hansen, T. Hansen, V. Haroutunian, A. M. Hartmann, F. A. Henskens, S. Herms, J. N. Hirschhorn, P. Hoffmann, A. Hofman, H. Huang, M. Ikeda, I. Joa, A. K. Kahler, R. S. Kahn, L. Kalaydjieva, J. Karjalainen, D. Kavanagh, M. C. Keller, B. J. Kelly, J. L. Kennedy, Y. Kim, J. A. Knowles, B. Konte, C. Laurent, P. Lee, S. H. Lee, S. E. Legge, B. Lerer, D. L. Levy, K. Y. Liang, J. Lieberman, J. Lonnqvist, C. M. Loughland, P. K. E. Magnusson, B. S. Maher, W. Maier, J. Mallet, M. Mattheisen, M. Mattingsdal, R. W. McCarley, C. McDonald, A. M. McIntosh, S. Meier, C. J. Meijer, I. Melle, R. I. Mesholam-Gately, A. Metspalu, P. T. Michie, L. Milani, V. Milanova, Y. Mokrab, D. W. Morris, B. Muller-Myhsok, K. C. Murphy, R. M. Murray, I. Myin-Germeys, I. Nenadic, D. A. Nertney, G. Nestadt, K. K. Nicodemus, L. Nisenbaum, A. Nordin, E. O’Callaghan, C. O’Dushlaine, S. Y. Oh, A. Olincy, L. Olsen, F. A. O’Neill, J. Van Os, C. Pantelis, G. N. Papadimitriou, E. Parkhomenko, M. T. Pato, T. Paunio, C. Psychosis Endophenotypes International, D. O. Perkins, T. H. Pers, O. Pietilainen, J. Pimm, A. J. Pocklington, J. Powell, A. Price, A. E. Pulver, S. M. Purcell, D. Quested, H. B. Rasmussen, A. Reichenberg, M. A. Reimers, A. L. Richards, J. L. Roffman, P. Roussos, D. M. Ruderfer, V. Salomaa, A. R. Sanders, A. Savitz, U. Schall, T. G. Schulze, S. G. Schwab, E. M. Scolnick, R. J. Scott, L. J. Seidman, J. Shi, J. M. Silverman, J. W. Smoller, E. Soderman, C. C. A. Spencer, E. A. Stahl, E. Strengman, J. Strohmaier, T. S. Stroup, J. Suvisaari, D. M. Svrakic, J. P. Szatkiewicz, S. Thirumalai, P. A. Tooney, J. Veijola, P. M. Visscher, J. Waddington, D. Walsh, B. T. Webb, M. Weiser, D. B. Wildenauer, N. M. Williams, S. Williams, S. H. Witt, A. R. Wolen, B. K. Wormley, N. R. Wray, J. Q. Wu, C. C. Zai, R. Adolfsson, O. A. Andreassen, D. H. R. Blackwood, E. Bramon, J. D. Buxbaum, S. Cichon, D. A. Collier, A. Corvin, M. J. Daly, A. Darvasi, E. Domenici, T. Esko, P. V. Gejman, M. Gill, H. Gurling, C. M. Hultman, N. Iwata, A. V. Jablensky, E. G. Jonsson, K. S. Kendler, G. Kirov, J. Knight, D. F. Levinson, Q. S. Li, S. A. McCarroll, A. McQuillin, J. L. Moran, B. J. Mowry, M. M. Nothen, R. A. Ophoff, M. J. Owen, A. Palotie, C. N. Pato, T. L. Petryshen, D. Posthuma, M. Rietschel, B. P. Riley, D. Rujescu, P. Sklar, D. St Clair, J. T. R. Walters, T. Werge, P. F. Sullivan, M. C. O’Donovan, S. W. Scherer, B. M. Neale, J. Sebat, Cnv, C. Schizophrenia Working Groups of the Psychiatric Genomics, Contribution of copy number variants to schizophrenia from a genome-wide study of 41,321 subjects. Nat Genet 49, 27–35 (2017).

7. G. Genovese, M. Fromer, E. A. Stahl, D. M. Ruderfer, K. Chambert, M. Landen, J. L. Moran, S. M. Purcell, P. Sklar, P. F. Sullivan, C. M. Hultman, S. A. McCarroll, Increased burden of ultra-rare protein-altering variants among 4,877 individuals with schizophrenia. Nature Neuroscience 19, 1433–1441 (2016).

8. T. Singh, M. I. Kurki, D. Curtis, S. M. Purcell, L. Crooks, J. McRae, J. Suvisaari, H. Chheda, D. Blackwood, G. Breen, O. Pietilainen, S. S. Gerety, M. Ayub, M. Blyth, T. Cole, D. Collier, E. L. Coomber, N. Craddock, M. J. Daly, J. Danesh, M. DiForti, A. Foster, N. B. Freimer, D. Geschwind, M. Johnstone, S. Joss, G. Kirov, J. Korkko, O. Kuismin, P. Holmans, C. M. Hultman, C. Iyegbe, J. Lonnqvist, M. Mannikko, S. A. McCarroll, P. McGuffin, A. M. McIntosh, A. McQuillin, J. S. Moilanen, C. Moore, R. M. Murray, R. Newbury-Ecob, W. Ouwehand, T. Paunio, E. Prigmore, E. Rees, D. Roberts, J. Sambrook, P. Sklar, D. St Clair, J. Veijola, J. T. Walters, H. Williams, S. Swedish Schizophrenia, I. Study, D. D. D. Study, U. K. Consortium, P. F. Sullivan, M. E. Hurles, M. C. O’Donovan, A. Palotie, M. J. Owen, J. C. Barrett, Rare loss-of-function variants in SETD1A are associated with schizophrenia and developmental disorders. Nat Neurosci 19, 571–577 (2016).

9. E. A. Boyle, Y. I. Li, J. K. Pritchard, An Expanded View of Complex Traits: From Polygenic to Omnigenic. Cell 169, 1177–1186 (2017).

10. E. Mondragon, L. J. Maher, 3rd, Anti-Transcription Factor RNA Aptamers as Potential Therapeutics. Nucleic Acid Ther 26, 29–43 (2016).

11. B. F. Corbett, J. C. You, X. Zhang, M. S. Pyfer, U. Tosi, D. M. Iascone, I. Petrof, A. Hazra, C. H. Fu, G. S. Stephens, A. A. Ashok, S. Aschmies, L. Zhao, E. J. Nestler, J. Chin, DeltaFosB Regulates Gene Expression and Cognitive Dysfunction in a Mouse Model of Alzheimer’s Disease. Cell Rep 20, 344–355 (2017).

12. M. Lizio, J. Harshbarger, H. Shimoji, J. Severin, T. Kasukawa, S. Sahin, I. Abugessaisa, S. Fukuda, F. Hori, S. Ishikawa-Kato, C. J. Mungall, E. Arner, J. K. Baillie, N. Bertin, H. Bono, M. de Hoon, A. D. Diehl, E. Dimont, T. C. Freeman, K. Fujieda, W. Hide, R. Kaliyaperumal, T. Katayama, T. Lassmann, T. F. Meehan, K. Nishikata, H. Ono, M. Rehli, A. Sandelin, E. A. Schultes, P. A. t Hoen, Z. Tatum, M. Thompson, T. Toyoda, D. W. Wright, C. O. Daub, M. Itoh, P. Carninci, Y. Hayashizaki, A. R. Forrest, H. Kawaji, F. consortium, Gateways to the FANTOM5 promoter level mammalian expression atlas. Genome Biol 16, 22 (2015).

13. J. M. Vaquerizas, S. K. Kummerfeld, S. A. Teichmann, N. M. Luscombe, A census of human transcription factors: function, expression and evolution. Nat Rev Genet 10, 252–263 (2009).

14. H. Han, H. Shim, D. Shin, J. E. Shim, Y. Ko, J. Shin, H. Kim, A. Cho, E. Kim, T. Lee, H. Kim, K. Kim, S. Yang, D. Bae, A. Yun, S. Kim, C. Y. Kim, H. J. Cho, B. Kang, S. Shin, I. Lee, TRRUST: a reference database of human transcriptional regulatory interactions. Sci Rep 5, 11432 (2015).

15. I. K. Jordan, L. Marino-Ramirez, Y. I. Wolf, E. V. Koonin, Conservation and coevolution in the scale-free human gene coexpression network. Mol Biol Evol 21, 2058–2070 (2004).

16. K. Basso, A. A. Margolin, G. Stolovitzky, U. Klein, R. Dalla-Favera, A. Califano, Reverse engineering of regulatory networks in human B cells. Nat Genet 37, 382–390 (2005).

17. A. A. Margolin, K. Wang, W. K. Lim, M. Kustagi, I. Nemenman, A. Califano, Reverse engineering cellular networks. Nat Protoc 1, 662–671 (2006).

18. M. J. Alvarez, Y. Shen, F. M. Giorgi, A. Lachmann, B. B. Ding, B. H. Ye, A. Califano, Functional characterization of somatic mutations in cancer using network-based inference of protein activity. Nature Genetics 48, 838–+ (2016).

19. L. Brichta, W. Shin, V. Jackson-Lewis, J. Blesa, E. L. Yap, Z. Walker, J. Zhang, J. P. Roussarie, M. J. Alvarez, A. Califano, S. Przedborski, P. Greengard, Identification of neurodegenerative factors using translatome-regulatory network analysis. Nat Neurosci 18, 1325–+ (2015).

20. V. Haroutunian, P. Katsel, S. Dracheva, D. G. Stewart, K. L. Davis, Variations in oligodendrocyte-related gene expression across multiple cortical regions: implications for the pathophysiology of schizophrenia. Int J Neuropsychopharmacol 10, 565–573 (2007).

21. D. Arion, S. Horvath, D. A. Lewis, K. Mirnics, Infragranular gene expression disturbances in the prefrontal cortex in schizophrenia: signature of altered neural development? Neurobiol Dis 37, 738–746 (2010).

22. A. L. Guillozet-Bongaarts, T. M. Hyde, R. A. Dalley, M. J. Hawrylycz, A. Henry, P. R. Hof, J. Hohmann, A. R. Jones, C. L. Kuan, J. Royall, E. Shen, B. Swanson, H. Zeng, J. E. Kleinman, Altered gene expression in the dorsolateral prefrontal cortex of individuals with schizophrenia. Mol Psychiatry 19, 478–485 (2014).

23. M. Fromer, P. Roussos, S. K. Sieberts, J. S. Johnson, D. H. Kavanagh, T. M. Perumal, D. M. Ruderfer, E. C. Oh, A. Topol, H. R. Shah, L. L. Klei, R. Kramer, D. Pinto, Z. H. Gumus, A. E. Cicek, K. K. Dang, A. Browne, C. Lu, L. Xie, B. Readhead, E. A. Stahl, J. Xiao, M. Parvizi, T. Hamamsy, J. F. Fullard, Y. C. Wang, M. C. Mahajan, J. M. Derry, J. T. Dudley, S. E. Hemby, B. A. Logsdon, K. Talbot, T. Raj, D. A. Bennett, P. L. De Jager, J. Zhu, B. Zhang, P. F. Sullivan, A. Chess, S. M. Purcell, L. A. Shinobu, L. M. Mangravite, H. Toyoshiba, R. E. Gur, C. G. Hahn, D. A. Lewis, V. Haroutunian, M. A. Peters, B. K. Lipska, J. D. Buxbaum, E. E. Schadt, K. Hirai, K. Roeder, K. J. Brennand, N. Katsanis, E. Domenici, B. Devlin, P. Sklar, Gene expression elucidates functional impact of polygenic risk for schizophrenia. Nat Neurosci 19, 1442–1453 (2016).

24. O. V. Evgrafov, C. Armoskus, B. B. Wrobel, V. N. Spitsyna, T. Souaiaia, J. S. Herstein, C. P. Walker, J. D. Nguyen, A. Camarena, J. R. Weitz, J. M. H. Kim, E. Lopez Duarte, K. Wang, G. M. Simpson, J. L. Sobell, H. Medeiros, M. T. Pato, C. N. Pato, J. A. Knowles, Gene expression in patient-derived neural progenitors provide insights into neurodevelopmental aspects of schizophrenia. bioRxiv, (2017).

25. C. Lefebvre, P. Rajbhandari, M. J. Alvarez, P. Bandaru, W. K. Lim, M. Sato, K. Wang, P. Sumazin, M. Kustagi, B. C. Bisikirska, K. Basso, P. Beltrao, N. Krogan, J. Gautier, R. Dalla-Favera, A. Califano, A human B-cell interactome identifies MYB and FOXM1 as master regulators of proliferation in germinal centers. Mol Syst Biol 6, (2010).

26. K. Yang, A. Sawa, Open Sesame: Open Chromatin Regions Shed Light onto Non-coding Risk Variants. Cell Stem Cell 21, 285–287 (2017).

27. B. B. Lake, S. Chen, B. C. Sos, J. Fan, G. E. Kaeser, Y. C. Yung, T. E. Duong, D. Gao, J. Chun, P. V. Kharchenko, K. Zhang, Integrative single-cell analysis of transcriptional and epigenetic states in the human adult brain. Nat Biotechnol 36, 70–80 (2018).

28. S. Steinberg, S. de Jong, C. Irish Schizophrenia Genomics, O. A. Andreassen, T. Werge, A. D. Borglum, O. Mors, P. B. Mortensen, O. Gustafsson, J. Costas, O. P. Pietilainen, D. Demontis, S. Papiol, J. Huttenlocher, M. Mattheisen, R. Breuer, E. Vassos, I. Giegling, G. Fraser, N. Walker, A. Tuulio-Henriksson, J. Suvisaari, J. Lonnqvist, T. Paunio, I. Agartz, I. Melle, S. Djurovic, E. Strengman, Group, G. Jurgens, B. Glenthoj, L. Terenius, D. M. Hougaard, T. Orntoft, C. Wiuf, M. Didriksen, M. V. Hollegaard, M. Nordentoft, R. van Winkel, G. Kenis, L. Abramova, V. Kaleda, M. Arrojo, J. Sanjuan, C. Arango, S. Sperling, M. Rossner, M. Ribolsi, V. Magni, A. Siracusano, C. Christiansen, L. A. Kiemeney, J. Veldink, L. van den Berg, A. Ingason, P. Muglia, R. Murray, M. M. Nothen, E. Sigurdsson, H. Petursson, U. Thorsteinsdottir, A. Kong, I. A. Rubino, M. De Hert, J. M. Rethelyi, I. Bitter, E. G. Jonsson, V. Golimbet, A. Carracedo, H. Ehrenreich, N. Craddock, M. J. Owen, M. C. O’Donovan, C. Wellcome Trust Case Control, M. Ruggeri, S. Tosato, L. Peltonen, R. A. Ophoff, D. A. Collier, D. St Clair, M. Rietschel, S. Cichon, H. Stefansson, D. Rujescu, K. Stefansson, Common variants at VRK2 and TCF4 conferring risk of schizophrenia. Hum Mol Genet 20, 4076–4081 (2011).

29. M. Kharbanda, K. Kannike, A. Lampe, J. Berg, T. Timmusk, M. Sepp, Partial deletion of TCF4 in three generation family with non-syndromic intellectual disability, without features of Pitt-Hopkins syndrome. Eur J Med Genet 59, 310–314 (2016).

30. M. P. Forrest, M. J. Hill, A. J. Quantock, E. Martin-Rendon, D. J. Blake, The emerging roles of TCF4 in disease and development. Trends Mol Med 20, 322–331 (2014).

31. J. Wang, D. Duncan, Z. Shi, B. Zhang, WEB-based GEne SeT AnaLysis Toolkit (WebGestalt): update 2013. Nucleic Acids Res 41, W77–83 (2013).

32. A. Mathelier, O. Fornes, D. J. Arenillas, C. Y. Chen, G. Denay, J. Lee, W. Shi, C. Shyr, G. Tan, R. Worsley-Hunt, A. W. Zhang, F. Parcy, B. Lenhard, A. Sandelin, W. W. Wasserman, JASPAR 2016: a major expansion and update of the open-access database of transcription factor binding profiles. Nucleic Acids Res 44, D110–115 (2016).

33. R. I. Sherwood, T. Hashimoto, C. W. O’Donnell, S. Lewis, A. A. Barkal, J. P. van Hoff, V. Karun, T. Jaakkola, D. K. Gifford, Discovery of directional and nondirectional pioneer transcription factors by modeling DNase profile magnitude and shape. Nat Biotechnol 32, 171–178 (2014).

34. M. P. Forrest, H. Zhang, W. Moy, H. McGowan, C. Leites, L. E. Dionisio, Z. Xu, J. Shi, A. R. Sanders, W. J. Greenleaf, C. A. Cowan, Z. P. Pang, P. V. Gejman, P. Penzes, J. Duan, Open Chromatin Profiling in hiPSC-Derived Neurons Prioritizes Functional Noncoding Psychiatric Risk Variants and Highlights Neurodevelopmental Loci. Cell Stem Cell 21, 305–318 e308 (2017).

35. S. Zhang, W. Moy, H. Zhang, C. Leites, H. McGowan, J. Shi, A. R. Sanders, Z. P. Pang, P. V. Gejman, J. Duan, Open chromatin dynamics reveals stage-specific transcriptional networks in hiPSC-based neurodevelopmental model. Stem Cell Res 29, 88–98 (2018).

36. M. P. Forrest, M. J. Hill, D. H. Kavanagh, K. E. Tansey, A. J. Waite, D. J. Blake, The Psychiatric Risk Gene Transcription Factor 4 (TCF4) Regulates Neurodevelopmental Pathways Associated With Schizophrenia, Autism, and Intellectual Disability. Schizophrenia Bulletin, sbx164–sbx164 (2017).

37. H. Xia, F. M. Jahr, N. K. Kim, L. Xie, A. A. Shabalin, J. Bryois, D. H. Sweet, M. M. Kronfol, P. Palasuberniam, M. McRae, B. P. Riley, P. F. Sullivan, E. J. van den Oord, J. L. McClay, Building a schizophrenia genetic network: transcription factor 4 regulates genes involved in neuronal development and schizophrenia risk. Hum Mol Genet 27, 3246–3256 (2018).

38. M. P. Forrest, A. J. Waite, E. Martin-Rendon, D. J. Blake, Knockdown of human TCF4 affects multiple signaling pathways involved in cell survival, epithelial to mesenchymal transition and neuronal differentiation. PLoS One 8, e73169 (2013).

39. A. J. Kennedy, E. J. Rahn, B. S. Paulukaitis, K. E. Savell, H. B. Kordasiewicz, J. Wang, J. W. Lewis, J. Posey, S. K. Strange, M. C. Guzman-Karlsson, S. E. Phillips, K. Decker, S. T. Motley, E. E. Swayze, D. J. Ecker, T. P. Michael, J. J. Day, J. D. Sweatt, Tcf4 Regulates Synaptic Plasticity, DNA Methylation, and Memory Function. Cell Rep 16, 2666–2685 (2016).

40. M. D. Rannals, G. R. Hamersky, S. C. Page, M. N. Campbell, A. Briley, R. A. Gallo, B. N. Phan, T. M. Hyde, J. E. Kleinman, J. H. Shin, A. E. Jaffe, D. R. Weinberger, B. J. Maher, Psychiatric Risk Gene Transcription Factor 4 Regulates Intrinsic Excitability of Prefrontal Neurons via Repression of SCN10a and KCNQ1. Neuron 90, 43–55 (2016).

41. A. Devor, O. A. Andreassen, Y. Wang, T. Maki-Marttunen, O. B. Smeland, C. C. Fan, A. J. Schork, D. Holland, W. K. Thompson, A. Witoelar, C. H. Chen, R. S. Desikan, L. K. McEvoy, S. Djurovic, P. Greengard, P. Svenningsson, G. T. Einevoll, A. M. Dale, Genetic evidence for role of integration of fast and slow neurotransmission in schizophrenia. Mol Psychiatry 22, 792–801 (2017).

42. A. F. Pardinas, P. Holmans, A. J. Pocklington, V. Escott-Price, S. Ripke, N. Carrera, S. E. Legge, S. Bishop, D. Cameron, M. L. Hamshere, J. Han, L. Hubbard, A. Lynham, K. Mantripragada, E. Rees, J. H. MacCabe, S. A. McCarroll, B. T. Baune, G. Breen, E. M. Byrne, U. Dannlowski, T. C. Eley, C. Hayward, N. G. Martin, A. M. McIntosh, R. Plomin, D. J. Porteous, N. R. Wray, A. Caballero, D. H. Geschwind, L. M. Huckins, D. M. Ruderfer, E. Santiago, P. Sklar, E. A. Stahl, H. Won, E. Agerbo, T. D. Als, O. A. Andreassen, M. Baekvad-Hansen, P. B. Mortensen, C. B. Pedersen, A. D. Borglum, J. Bybjerg-Grauholm, S. Djurovic, N. Durmishi, M. G. Pedersen, V. Golimbet, J. Grove, D. M. Hougaard, M. Mattheisen, E. Molden, O. Mors, M. Nordentoft, M. Pejovic-Milovancevic, E. Sigurdsson, T. Silagadze, C. S. Hansen, K. Stefansson, H. Stefansson, S. Steinberg, S. Tosato, T. Werge, G. Consortium, C. Consortium, D. A. Collier, D. Rujescu, G. Kirov, M. J. Owen, M. C. O’Donovan, J. T. R. Walters, G. Consortium, C. Consortium, G. Consortium, C. Consortium, Common schizophrenia alleles are enriched in mutation-intolerant genes and in regions under strong background selection. Nat Genet 50, 381–389 (2018).

43. P. Langfelder, S. Horvath, WGCNA: an R package for weighted correlation network analysis. BMC Bioinformatics 9, 559 (2008).

44. A. S. Boyajyan, S. A. Atshemyan, R. V. Zakharyan, Association of schizophrenia with variants of genes that encode transcription factors. Mol Biol+ 49, 875–880 (2015).

45. E. S. Emamian, AKT/GSK3 signaling pathway and schizophrenia. Front Mol Neurosci 5, 33 (2012).

46. A. Y. Guo, J. Sun, P. Jia, Z. Zhao, A novel microRNA and transcription factor mediated regulatory network in schizophrenia. BMC Syst Biol 4, 10 (2010).

47. E. Masliah, W. Dumaop, D. Galasko, P. Desplats, Distinctive patterns of DNA methylation associated with Parkinson disease: identification of concordant epigenetic changes in brain and peripheral blood leukocytes. Epigenetics 8, 1030–1038 (2013).

48. A. O. M. Bennett, Stress and anxiety in schizophrenia and depression: glucocorticoids, corticotropin-releasing hormone and synapse regression. Aust N Z J Psychiatry 42, 995–1002 (2008).

49. P. R. Maycox, F. Kelly, A. Taylor, S. Bates, J. Reid, R. Logendra, M. R. Barnes, C. Larminie, N. Jones, M. Lennon, C. Davies, J. J. Hagan, C. A. Scorer, C. Angelinetta, M. T. Akbar, S. Hirsch, A. M. Mortimer, T. R. Barnes, J. de Belleroche, Analysis of gene expression in two large schizophrenia cohorts identifies multiple changes associated with nerve terminal function. Mol Psychiatry 14, 1083–1094 (2009).

50. N. Sugo, H. Oshiro, M. Takemura, T. Kobayashi, Y. Kohno, N. Uesaka, W. J. Song, N. Yamamoto, Nucleocytoplasmic translocation of HDAC9 regulates gene expression and dendritic growth in developing cortical neurons. Eur J Neurosci 31, 1521–1532 (2010).

51. J. P. Lopez-Atalaya, S. Ito, L. M. Valor, E. Benito, A. Barco, Genomic targets, and histone acetylation and gene expression profiling of neural HDAC inhibition. Nucleic Acids Res 41, 8072–8084 (2013).

52. Y. Fan, G. Abrahamsen, R. Mills, C. C. Calderon, J. Y. Tee, L. Leyton, W. Murrell, J. Cooper-White, J. J. McGrath, A. Mackay-Sim, Focal adhesion dynamics are altered in schizophrenia. Biol Psychiatry 74, 418–426 (2013).

53. L. J. Carithers, K. Ardlie, M. Barcus, P. A. Branton, A. Britton, S. A. Buia, C. C. Compton, D. S. DeLuca, J. Peter-Demchok, E. T. Gelfand, P. Guan, G. E. Korzeniewski, N. C. Lockhart, C. A. Rabiner, A. K. Rao, K. L. Robinson, N. V. Roche, S. J. Sawyer, A. V. Segre, C. E. Shive, A. M. Smith, L. H. Sobin, A. H. Undale, K. M. Valentino, J. Vaught, T. R. Young, H. M. Moore, G. T. Consortium, A Novel Approach to High-Quality Postmortem Tissue Procurement: The GTEx Project. Biopreserv Biobank 13, 311–319 (2015).

54. H. Ding, E. F. Douglass, Jr., A. M. Sonabend, A. Mela, S. Bose, C. Gonzalez, P. D. Canoll, P. A. Sims, M. J. Alvarez, A. Califano, Quantitative assessment of protein activity in orphan tissues and single cells using the metaVIPER algorithm. Nat Commun 9, 1471 (2018).

55. D. Wang, S. Liu, J. Warrell, H. Won, X. Shi, F. C. P. Navarro, D. Clarke, M. Gu, P. Emani, Y. T. Yang, M. Xu, M. J. Gandal, S. Lou, J. Zhang, J. J. Park, C. Yan, S. K. Rhie, K. Manakongtreecheep, H. Zhou, A. Nathan, M. Peters, E. Mattei, D. Fitzgerald, T. Brunetti, J. Moore, Y. Jiang, K. Girdhar, G. E. Hoffman, S. Kalayci, Z. H. Gumus, G. E. Crawford, E. C. Psych, P. Roussos, S. Akbarian, A. E. Jaffe, K. P. White, Z. Weng, N. Sestan, D. H. Geschwind, J. A. Knowles, M. B. Gerstein, Comprehensive functional genomic resource and integrative model for the human brain. Science 362, (2018).

56. H. K. Finucane, Y. A. Reshef, V. Anttila, K. Slowikowski, A. Gusev, A. Byrnes, S. Gazal, P. R. Loh, C. Lareau, N. Shoresh, G. Genovese, A. Saunders, E. Macosko, S. Pollack, C. Brainstorm, J. R. B. Perry, J. D. Buenrostro, B. E. Bernstein, S. Raychaudhuri, S. McCarroll, B. M. Neale, A. L. Price, Heritability enrichment of specifically expressed genes identifies disease-relevant tissues and cell types. Nat Genet 50, 621–629 (2018).

57. A. Yates, W. Akanni, M. R. Amode, D. Barrell, K. Billis, D. Carvalho-Silva, C. Cummins, P. Clapham, S. Fitzgerald, L. Gil, C. G. Giron, L. Gordon, T. Hourlier, S. E. Hunt, S. H. Janacek, N. Johnson, T. Juettemann, S. Keenan, I. Lavidas, F. J. Martin, T. Maurel, W. McLaren, D. N. Murphy, R. Nag, M. Nuhn, A. Parker, M. Patricio, M. Pignatelli, M. Rahtz, H. S. Riat, D. Sheppard, K. Taylor, A. Thormann, A. Vullo, S. P. Wilder, A. Zadissa, E. Birney, J. Harrow, M. Muffato, E. Perry, M. Ruffier, G. Spudich, S. J. Trevanion, F. Cunningham, B. L. Aken, D. R. Zerbino, P. Flicek, Ensembl 2016. Nucleic Acids Res 44, D710–716 (2016).

58. J. Shi, D. F. Levinson, J. Duan, A. R. Sanders, Y. Zheng, I. Pe’er, F. Dudbridge, P. A. Holmans, A. S. Whittemore, B. J. Mowry, A. Olincy, F. Amin, C. R. Cloninger, J. M. Silverman, N. G. Buccola, W. F. Byerley, D. W. Black, R. R. Crowe, J. R. Oksenberg, D. B. Mirel, K. S. Kendler, R. Freedman, P. V. Gejman, Common variants on chromosome 6p22.1 are associated with schizophrenia. Nature 460, 753–757 (2009).

59. A. Dobin, C. A. Davis, F. Schlesinger, J. Drenkow, C. Zaleski, S. Jha, P. Batut, M. Chaisson, T. R. Gingeras, STAR: ultrafast universal RNA-seq aligner. Bioinformatics 29, 15–21 (2013).

60. M. D. Robinson, D. J. McCarthy, G. K. Smyth, edgeR: a Bioconductor package for differential expression analysis of digital gene expression data. Bioinformatics 26, 139–140 (2010).

